# Structural insights into the mechanism of the human soluble guanylate cyclase

**DOI:** 10.1101/731679

**Authors:** Yunlu Kang, Rui Liu, Jing-Xiang Wu, Lei Chen

**Author notes:** These authors contribute equally to this work (Yunlu Kang, Rui Liu). Correspondence and Lead Contact: Lei Chen.

## Abstract

Soluble guanylate cyclase (sGC) is the primary nitric oxide (NO) sensor. It plays a central role in NO signaling and is implicated in many essential physiological processes and disease conditions. The binding of NO leads to a significant boost in sGC enzymatic activity. However, the mechanism of NO activation remains incompletely understood. Here, we report the cryo-electron microscopy structures of the human sGC α1β1 heterodimer in different functional states. These structures revealed that the transducer module bridges the NO sensor module and the catalytic module. NO binding to the β1 H-NOX domain triggers the structural rearrangement of the sensor module and the bending-straightening conformational switch of the transducer module. The resulting movement of the N-termini of the catalytic domains drives the structural changes within the catalytic module, which in turn boost sGC enzymatic activity. These observations indicate the structural framework for the mechanism of sGC activation induced by NO binding.

## Introduction

Nitric oxide (NO) is a unique gaseous signaling molecule involved in many important physiological processes, such as vasodilatation, neurotransmission, platelet aggregation, immunity, cell proliferation, and mitochondrial respiration (Hollenberg and Cinel, 2009; Horst and Marletta, 2018). The dysregulation of NO signaling has been linked to cardiovascular disease, sepsis, acute lung injury, and multiple organ failure (Farah et al., 2018; Hollenberg and Cinel, 2009; Luiking et al., 2010). NO signaling is initiated by the activation of NO synthase (NOS), which generates NO in response to physiologic stimuli. NO readily permeates target cell membranes, and after diffusing across the membrane, it binds and activates soluble guanylate cyclase (sGC), the primary NO acceptor. sGC catalyzes the cyclization reaction of GTP to generate inorganic pyrophosphate and the secondary messenger cyclic guanosine monophosphate (cGMP). cGMP then acts on the downstream effectors, including cGMP-regulated protein kinases, phosphodiesterases, and ion channels, to regulate physiological processes in the cell (Derbyshire and Marletta, 2012). Therefore, sGC acts as both a sensor and an amplifier in NO signaling. Genetic mutation of sGC in humans is associated with coronary artery disease (Deloukas et al., 2013), moyamoya disease, achalasia, and hypertension (Herve et al., 2014; Wallace et al., 2016), and it is a validated drug target for the treatment of pulmonary hypertension and chronic heart failure. The NO donor nitroglycerin has been widely used for centuries to alleviate angina pectoris, and the sGC activator riociguat has been approved for the treatment of pulmonary hypertension. sGC activators also have therapeutic potential in fibrotic diseases, systemic sclerosis, chronic kidney diseases, neuroprotection, dementia, and sickle cell disease (Sandner, 2018).

sGC is a heterodimeric protein complex composed of one α and one β subunit. In humans, the α subunit can be either α1 or α2 and the β subunit can be either β1 or β2. The α1 and β1 subunits are widely expressed in many tissues, while the expression of α2 and β2 subunits is tissue-specific (Derbyshire and Marletta, 2012; Koesling et al., 2016). The α and β subunits have sequence homology to some extent and are similarly organized into modular domains, including an N-terminal Heme-Nitric oxide and Oxygen binding (H-NOX) domain, a Per/Arnt/Sim (PAS) domain, a Coiled-Coil (CC) domain, and a C-terminal catalytic domain. The PAS and CC domains mediate protein-protein interactions, and the catalytic domain is responsible for enzymatic activity (Derbyshire and Marletta, 2012; Montfort et al., 2017). The H-NOX domain of the β subunit contains a ferrous b-type heme prosthetic group that facilitates the high affinity binding of NO (Derbyshire and Marletta, 2012; Montfort et al., 2017). Under pathological conditions or oxidative stress, the ferrous heme can be oxidized to ferric heme (Dasgupta et al., 2015), and heme-oxidized sGC has a poor response to NO, as evidenced by the markedly decreased NO-activation (Zhao et al., 2000).

To date, several structures of isolated sGC domains have been solved by X-ray crystallography or NMR. These structures include human β1 H-NOX, *Manduca sexta* α PAS domain (Purohit et al., 2013), human β1 CC domain (Ma et al., 2010), and human α1, β1 catalytic domain heterodimer (Allerston et al., 2013; Seeger et al., 2014). Recent negative stain electron microscope (EM) studies revealed the general shape of the full-length mammalian sGC at a resolution of 25-40 Å (Campbell et al., 2014), and hydrogen deuterium exchange experiments mapped NO-induced structural changes onto the primary sequence of the full-length sGC (Underbakke et al., 2014). Despite these pioneering structural works, the allosteric mechanism underlying the activation of the distal catalytic domain in response to NO binding to the β H-NOX domain remains unclear at the atomic level, mainly due to the lack of high-resolution structural information on intact sGC in different functional states. Here, we used cryo-EM to determine the structure of the human α1β1 sGC holoenzyme in both inactive and NO-activated states at a resolution of 3.9 Å and 3.8 Å, respectively. We also obtained a 6.8 Å resolution cryo-EM map of the constitutively active β1 H105C mutant. These structures uncover not only the detailed domain-domain interfaces, but also the activation mechanism of human sGC.

## Results and Discussions

### Structure determination

We found that sGC composed of human α1 and β1 can be expressed and purified to apparent homogeneity (Figures 1A, S1A and S1B) (Lee et al., 2000). sGC protein composed of an α1 and a β1 subunit is the most predominant isoform, and it has been widely used as a model protein to elucidate the biochemical, biophysical, and structural properties of mammalian sGC (Derbyshire and Marletta, 2012). Purified human α1β1 sGC showed characteristic UV-Vis spectrum of sGC with ferrous heme bound (Figure S1C) (Lee et al., 2000) and robust NO-activated GTP cyclase activity (Figure 1B), indicating that the prosthetic ferrous heme group was properly incorporated into the protein. In contrast, the β1 H105C mutant, in which the heme group is unable to bind, showed a constitutively high basal activity and was insensitive to NO activation (Figure 1B) (Martin et al., 2003). We prepared a heme-unliganded sGC sample, in which no exogenous ligand was supplemented, and then supplemented different small molecules to stabilize the purified sGC protein into functionally distinct states. The compound NS2028 has been reported to efficiently oxidize the Fe (II) in sGC to Fe (III) (Olesen et al., 1998). Indeed, we found that incubating sGC with NS2028 almost completely shifted the Soret peak from 431 to 392 nm (Figure S1C). Therefore, we incubated purified sGC with NS2028, Mg^2+^ ions, and substrate GTPγS (Brandwein et al., 1982) to obtain the heme-oxidized state. To achieve the NO-activated state, we supplemented the purified protein with excess NO donor DEA NONOate (Artz et al., 2001), Mg^2+^ ions, and noncyclizable substrate analogue GMPCPP (Cary et al., 2005).

**Figure 1.**
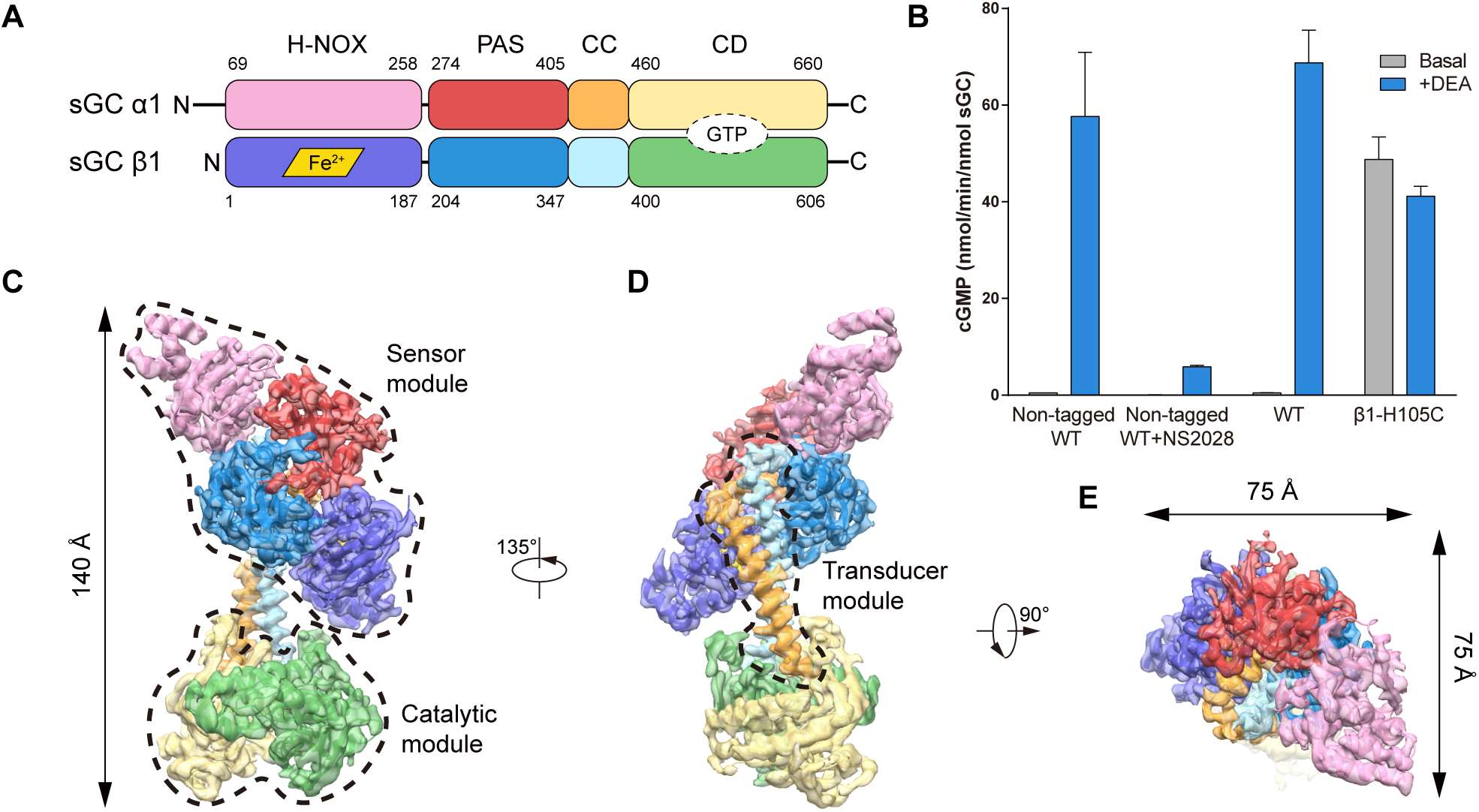
Structure of sGC in the inactive state. (A) Domain organization of the human α1β1 sGC heterodimer: Heme-Nitric oxide and Oxygen binding (H-NOX) domain, Per/Arnt/Sim (PAS) domain, Coiled-Coil (CC) domain and Catalytic Domain (CD). The approximate boundaries of the domains are listed as amino acid numbers. The heme cofactor and GTP substrate binding site are shown as a rhombus and an oval, respectively. (B) End-point activity assay of the purified non-tagged wild type sGC protein sample for cryo-EM, compared with that of NS2028 treated non-tagged wild type sGC, wild type sGC with CGFP tag (WT) and β1 H105C mutant with CGFP tag (Data are shown as means ± standard error, n = 3). (C) The side view of the cryo-EM map of sGC in the inactive (heme-oxidized) state. The colors of each individual domain are the same as in (A). The atomic model is shown as a cartoon inside the transparent electron density surface. The approximate boundaries of the sensor module and the catalytic module are shown in dashed lines. (D) A 135° rotated view compared to (B). The approximate boundary of the transducer module is shown in dashed lines. (E) A 90° rotated top view compared to (C).

The heme-unliganded, heme-oxidized, and NO-activated samples were subjected to single particle cryo-EM analysis (Figures S1-S4). During the 3D classification for each sample, we identified classes that had discernable secondary structural features (Figures S1F and S3C). These 3D classes were refined to overall resolution of 4 Å, 3.9 Å, and 3.8 Å to obtain the consensus maps for the heme-unliganded, heme-oxidized, and NO-activated states, respectively (Table S1). However, the two large lobes of sGC in the consensus maps showed blurry features, which were indicative of continuous conformational heterogeneities. Therefore, we divided the whole molecule into two bodies—the larger “N lobe” and the smaller “C lobe”—for further multibody refinement (Figures S1-S4) (Nakane et al., 2018), and the subsequent local map qualities were dramatically improved (Figures S2A-B and S3D-E). Analysis of the multibody refinements revealed monomodal particle distributions along the major eigenvectors given by the principal component analysis, indicating that the C lobe moved in a continuous fashion relative to the N lobe, probably due to the endogenous structural flexibility of the CC domains that connect the two lobes (Videos S1 and S2, Figures S2C-E and S4A-C). The two individual multibody maps were further aligned to the full consensus map and summed to generate the composite maps for model building and interpretation. The composite map qualities were sufficient to trace the main chain of most residues with the aid of available high-resolution homologous structures (Figures S2B, S2F, S3E, S4D, S5, S6, and Table S1). For the β1 H105C mutant sample, we added Mg^2+^ ions and the noncyclizable substrate analogue GMPCPP to the protein. The dataset of the β1 H105C mutant was processed as described above to obtain a reconstruction at 6.8 Å (Figures S4F-G). At this resolution, the overall shape and domain organization of β1 H105C mutant were found to be similar to that of the NO-activated state, which was low-passed to the same 6.8 Å resolution, with a real space correlation of 0.96. However, the heme density is missing in the β1 H-NOX domain of the β1 H105C mutant, as expected (Figure S4H).

### Structure of sGC in the inactive state

Both the heme-unliganded and the heme-oxidized sGC were in a contracted conformation, which is akin to the low resolution “bent” conformation observed previously in negative stain EM (Campbell et al., 2014) (Figures 1C-E, Video S3). We found that the overall structure of sGC in the heme-unliganded state (4 Å) is essentially the same as that in the heme-oxidized state (3.9 Å), with a RMSD of only 0.28 Å (Figure S7A). This was in accordance with the functional data, which showed that heme-unliganded state and heme-oxidized state have low activity (Figure 1B) (Zhao et al., 2000). Therefore, both of the structures were considered as the inactive state, and the 3.9 Å heme-oxidized state is used in further discussion of the inactive state due to its higher resolution and better map quality compared to the 4.0 Å heme-unliganded state. The structure of the inactive sGC occupies the 3D space of 140 Å × 75 Å × 75 Å (Figures 1C-E). The large N lobe is composed of α1 H-NOX, α1 PAS, β1 PAS, and β1 H-NOX domains. These domains are arranged in a pseudo two-fold symmetric manner, with the scaffolding PAS domains at the center and the H-NOX domains at the periphery (Figure 1C). These domains are essential for NO sensing and form the N-terminal “sensor” module of sGC (Figure 1C). The CC domains of both subunits form the “transducer” module that bridges the N-terminal sensor module and the C-terminal catalytic module (Figure 1D).

### Structure of the sGC sensor module in the inactive state

The atomic model of sGC in the inactive state allowed us to characterize the domain-domain interfaces in detail (Figures 2A-I). The N-terminal α1 H-NOX domain was previously proposed to share a common H-NOX fold, but the critical histidine residue for heme coordination is not conserved (Figures S5 and S6). Here, we show that the structure of the α1 H-NOX domain is similar to that of the β1 H-NOX domain. However, the N-terminal αA helix of the α1 H-NOX occupies a large portion of the pseudo-heme binding pocket and contributes to the formation of the H-NOX hydrophobic core (Figure S7B). Therefore, the α1 H-NOX domain does not retain the ability to bind heme under physiological conditions. The α1 H-NOX domain interacts with the adjacent PAS and β1 CC domains mainly through hydrophobic interactions (Figure 2B). βA, the αH-βD loop, and αF of the α1 H-NOX domain make contacts with the αK-βH loop of the α1 PAS domain. αA, αE, and αF of the α1 H-NOX domain interact with αL, the αL-αM loop, and αM of the β1 subunit (Figure 2B). Notably, there is no direct interaction observed between the α1 H-NOX and the β1 H-NOX domains, likely due to their distal spatial arrangement (Figure 2A).

**Figure 2.**
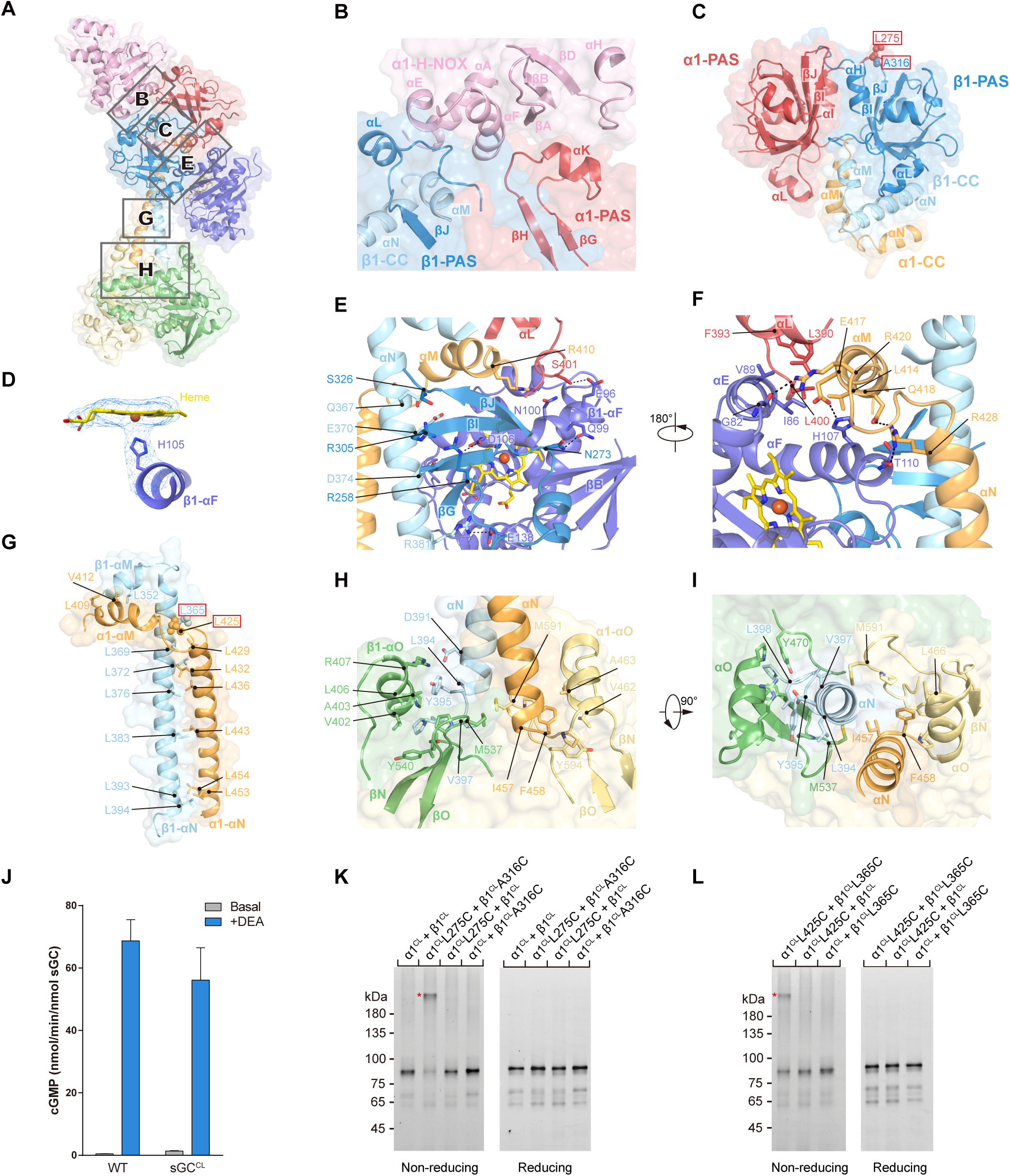
Domain-domain interfaces of sGC in the inactive state. (A) Side view of soluble guanylate cyclase in the inactive state, highlighting key interfaces (shown as gray rectangles). Each domain is colored the same as in Figure 1A. The surface of sGC is shown in transparency. (B) The interface between the α1 H-NOX domain and the PAS domains boxed in (A). (C) The interface between the PAS domains boxed in (A). The side chains of α1 L275 and β1 A316 which are in close proximity are shown as spheres. (D) The cryo-EM density map of the β1 H105-heme region in the inactive (heme-oxidized) sGC. The connectivity of densities between the heme and the histidine side chain is clearly identifiable. (E) The interface between β1 H-NOX and adjacent domains boxed in (A). (F) A 180° rotated view compared to (E). (G) The structure of the transducer module boxed in (A). The side chains of α1 L425 and β1 L365 which are in close proximity are shown as spheres. (H) The interface between the transducer module and the catalytic module boxed in (A). (I) A 90° rotated top view compared to (H). (J) End point activity of the cys-less construct compared to the wild type sGC with CGFP (sGC^CL^, α1^CL^+β1^CL^) (Data are shown as means ± standard error, n = 3). (K) SDS-PAGE of the *in vitro* disulfide bond cross-linking experiment of α1^CL^ L275 and β1^CL^ A316 mutants under reducing and non-reducing conditions. The in-gel GFP fluorescence of the α1 subunit is shown in black on white background. The position of cross-linked heterodimer is indicated with a red asterisk. (L) SDS-PAGE of the *in vitro* disulfide bond cross-linking experiment of α1^CL^ L425 and β1^CL^ L365 under reducing and non-reducing conditions. The position of cross-linked heterodimer is indicated with a red asterisk.

The structure of each PAS domain resembles that of the *Manduca sexta* sGC α subunit (PDB ID: 4GJ4) (Purohit et al., 2013) (Figures S7C-D). The α1 and β1 PAS domains are arranged into a pseudo-symmetric heterodimer (Figure 2C). Hydrophobic residues on αI, βI and βJ of the α1 PAS domain and αH, βI and βJ of the β1 PAS domain are packed into the stable hydrophobic core of the α1β1 PAS heterodimer (Figure 2C). In addition, αL of the PAS domain interacts with αM of the CC domain of the same subunit (Figure 2C). Interestingly, the β1 PAS domain interacts extensively with αN of the β1 CC domain. S326 on βJ hydrophilically interacts with Q367 of αN, and R305 on βI and R258 on βG interact electrostatically with E370 and D374 on αN, respectively (Figure 2E).

The H-NOX domain of the β1 subunit is the NO sensor of sGC (Derbyshire and Marletta, 2012). It shares common structural features with prokaryotic H-NOX domains (Plate and Marletta, 2013), which are composed of both N-terminal and C-terminal subdomains (Figure S7B). A heme molecule binds inside the β1 H-NOX domain, and its five coordinated Fe ion is tightly bound to H105 of αF, as evidenced by the strong connecting density between them (Figure 2D). β1 H-NOX extensively interacts with both the neighboring PAS heterodimers and the transducer module. On one side, I86 and V89 of αE hydrophobically interact with L390, F393, and L400 of the α1 PAS domain and L414 of the α1 CC domain, while G82, H107, and T110 of the β1 H-NOX domain form a complex hydrogen bonding network with R420, E417, and R428 of the α1 CC domain (Figure 2F). On the other side, E96, Q99, and N100 on αF of the β1 H-NOX domain form a polar interaction nexus with S401 of the α1 PAS domain, N273 of the β1 PAS domain, and R410 of the α1 CC domain (Figure 2E). In addition, E138 and D106 of the β1 H-NOX domain electrostatically interact with R381 of the β1 CC domain and R258 of the β1 PAS domain, respectively (Figure 2E).

### Structure of the sGC transducer module and the catalytic module in the inactive state

The polar interactions between the sensor module and the CC domains stabilize the transducer module in the “bent” conformation, in which both the α1 and β1 CC domains are broken into two short helixes (αM and αN) connected by a near 90° turn (Figure 2G). The α1 CC helix breaks at the forth turn (Figures 2G and S5) and the β1 CC helix breaks at the third turn (Figures 2G and S6). The αM and αN helixes of the α1 and β1 subunits still maintain two coiled-coil structures that are stabilized by the hydrophobic interactions of residues from both subunits (Figure 2G). The majority of these residues are leucines, which pack into structures reminiscent of leucine zippers (Figure 2G).

The C-terminal ends of the transducer module not only directly connect but also tightly interact with the catalytic module. I457 and F458 of the α1 CC insert into the hydrophobic pocket formed by A463, V462, L466, and Y594 of the α1 catalytic domain and M537 of the β1 catalytic domain (Figures 2H-I). In a similar fashion, L394, Y395, V397, and L398 of the β1 CC insert into the hydrophobic pocket formed by L406, A403, V402, Y540, and Y470 of the β1 catalytic domain and M591 of the α1 catalytic domain (Figures 2H-I). D391 of the β1 CC domain also electrostatically interacts with R407 of the β1 catalytic domain (Figure 2H). In the catalytic module, the two subunits are bound in a pseudo-symmetric manner (Figure S7E), consistent with previous observations of the isolated heterodimer (Seeger et al., 2014). However, the structure of the catalytic module in the inactive state of the full-length sGC is different from that of the isolated α1β1 catalytic domain heterodimer (Seeger et al., 2014). By aligning their β1 subunits, we found that the αQ helix of the α1 subunit rotates for about 13° (Figure S7F), indicating that the context of the full-length enzyme — especially the interactions between the transducer module and the catalytic module — was important for the structure of the catalytic module. When we compared the structure of the sGC catalytic module in the inactive state with that of adenylate cyclase in the active state (PDB ID: 1CJU) (Tesmer et al., 1999), we found that there are steric clashes between the adenosine group of the substrate and the α1 subunit of sGC and between the tri-phosphate group of the substrate and the β1 subunit of sGC (Figure S7G). This suggests that the structure of sGC in the inactive state is incompatible with substrate binding, which is in accordance with previous studies that showed that inactive sGC has a high *K*_m_ (Denninger et al., 2000).

To characterize and validate the domain-domain interactions observed in the inactive state, we generated a sGC mutant (named cys-less sGC, sGC^CL^) that has characteristic sGC enzymatic activity (Figure 2J) but fewer cysteines, and thus reduces background cross-linking (see Methods section). Cysteine mutations were introduced at spatially neighboring positions on both subunits. One pair (α1 L275C and β1 A316C) was located between the PAS domains, and the other pair (α1 L425C and β1 L365C) was located between the CC domains. We found that for each cysteine mutation pair, oxidative cross-linking only happened when the cysteine mutants were present in both subunit simultaneously (Figures 2K-L), which was consistent with our structural model of sGC in the inactive state.

### Structure of sGC in the NO-activated state

The NO-activated sGC has a dumbbell-shape structure in which the sensor module moves away from the catalytic module (Figure 3, Video S4). This is akin to the “extended” conformation previously observed by negative-stain EM (Campbell et al., 2014) and is dramatically different from the “bent” conformation of the inactive state. In addition, the overall structure of the constitutively active β1 H105C mutant shows a similar “extended” conformation in the absence of NO donor (Figure S4F). These structures suggest that this large overall conformational change is associated with the enhanced enzymatic activity of sGC but probably does not result from the covalent S-nitrosylation of sGC by NO, which can lead to sGC desensitization (Mayer et al., 2009; Sayed et al., 2007).

**Figure 3.**
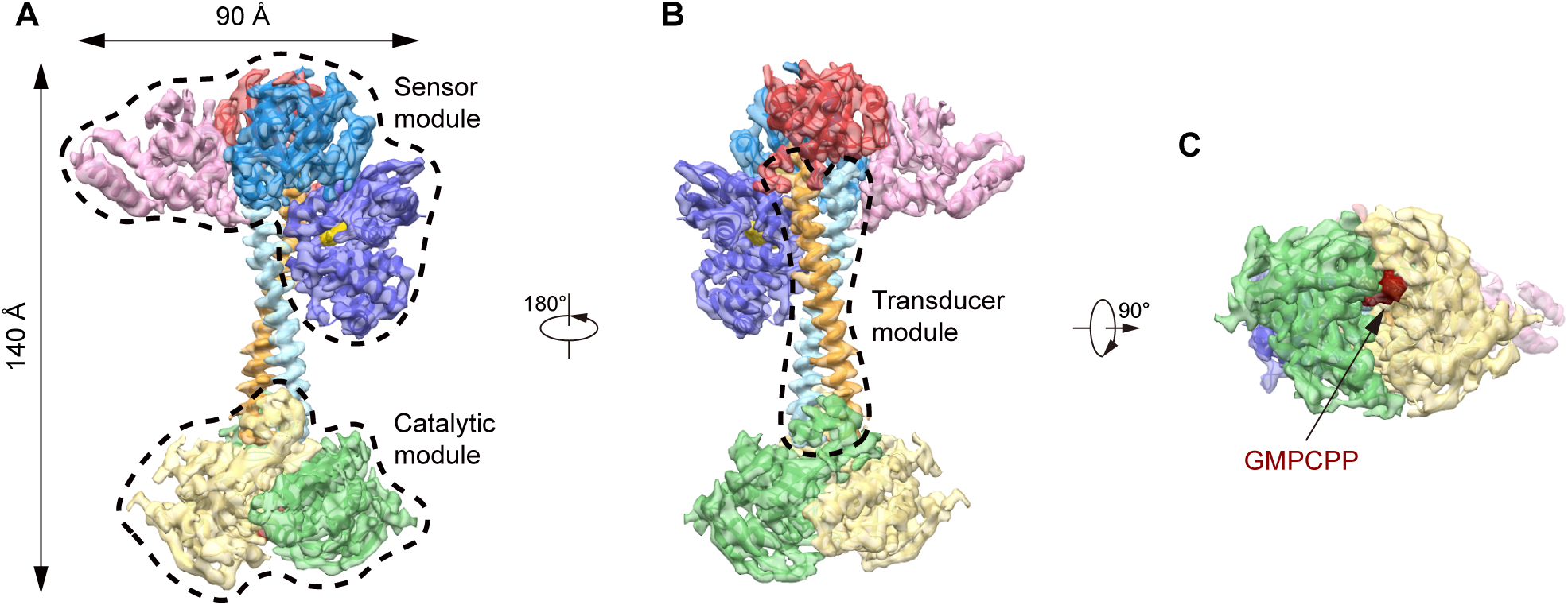
Structure of sGC in the NO-activated state. (A) Side view of the cryo-EM map of sGC in the NO-activated state. The colors of each individual domain are the same as in Figure 1A. The approximate boundaries of the sensor module and the catalytic module are shown in dashed lines. (B) A 180° rotated view compared to (A). The approximate boundary of the transducer module is shown in dashed lines. (C) A 90° rotated bottom view compared to (B). The density of GMPCPP is shown in dark red.

Despite the large overall conformational change, the general domain arrangement within each module in the NO-activated state is similar to that in the inactive state (Figures 1 and 3). In the electron density map of the NO-activated state, the H105-Fe bond of β1 H-NOX is cleaved, as evidenced by the clear separation between each density (Figure 4A). This suggests the current conformation corresponds to a NO-bound state, likely because excess NO donor DEA NONOate was added to the sample. However, we could not explicitly model the NO molecules or the heme deformation due to the limited resolution. The binding of NO induces a conformational change in β1 H-NOX in which the C-terminal subdomain rotates relative to the N-terminal subdomain (Figure S7H). As described in the inactive state structure, αE and αF of β1 H-NOX contribute to a large portion of the inter-domain interactions in the inactive state (Figures 2E-F). When αF (residues 96-107) of the β1 subunit was used as the reference to superimpose the structure of the NO-bound β1 H-NOX domain onto the structure of the inactive state, the Cα atom of N62 in the N-terminal subdomain has 4.6 Å displacement (Figure 4B). Furthermore, the NO-bound β1 H-NOX domain sterically clashes with the adjacent domains of the inactive state. Specifically, R40, I111, R116, and R121 of the NO-bound β1 H-NOX domain clash with Q422 and Q418 of the α1 CC domain, L380 of the β1 CC, and S270 of the β1 PAS domain of the inactive state, respectively (Figure 4B). This indicates that the inactive state structure is incompatible with the NO-bound β1 H-NOX domain and, therefore, a structural rearrangement is required to accommodate the conformational change of the β1 H-NOX domain upon NO binding. Indeed, we observed structural changes within the sensor module in which α1 H-NOX has a small downward movement while β1 H-NOX has a large rotational and translational movement (Figure 4C).

**Figure 4.**
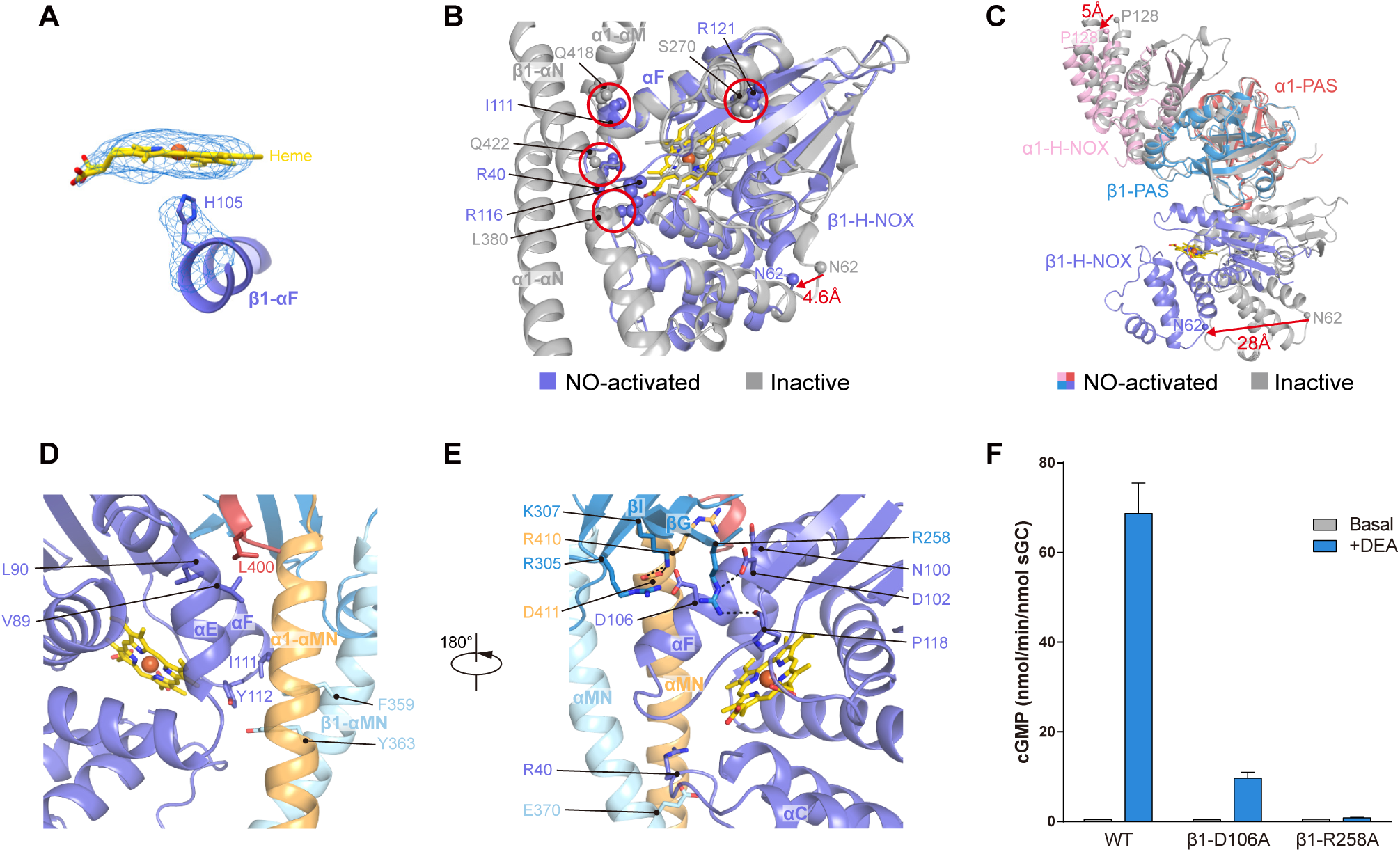
Domain-domain interfaces of sGC in the NO-activated state. (A) The cryo-EM density map of the β1 H105-heme region in the NO-activated sGC. The separation between the heme density and the histidine side chain density is clearly identifiable. (B) Superposition of the NO-bound β1 H-NOX domain structure (purple) onto the inactive state structure (gray) by alignment of the αF helixes. The steric clashes between the side chains of the NO-bound β1 H-NOX domain (purple sphere) and the side chains of the PAS and CC domains of the inactive state (gray sphere) are marked by red circles if their atom-to-atom distances are smaller than 2.2 Å. The arrow indicates the positional change of the Cα atoms of β1 N62 induced by NO binding. (C) Structural rearrangement of the sensor module. The structure of sGC in the NO-activated state (colored the same as Figure 1A) is superimposed onto the structure of sGC in the inactive state (gray) by aligning the PAS heterodimers. The arrows indicate the positional changes of the Cα atoms of α1 P128 and β1 N62 induced by NO binding. (D) The interface between β1 H-NOX and adjacent domains of sGC in the NO-activated state. (E) A 180° rotated view compared to (B). (F) End point activity assay of the β1 subunit mutants (Data are shown as means ± standard error, n = 3).

These conformational changes result in completely new interfaces between the NO-bound β1 H-NOX and adjacent domains (Figures 4D-E). Notably, V89 and L90 on αE of the β1 H-NOX domain hydrophobically interact with L400 of the α1 PAS domain, and I111 and Y112 in αF of the β1 H-NOX domain form hydrophobic interactions with F359 and Y363 of the β1 CC domain (Figure 4D). There are newly formed polar interactions as well: N100 and D106 on αF of the β1 H-NOX domain interact with R410 of the α1 CC domain and R305 of the β1 PAS domain, and K307 of the β1 PAS domain interacts with D411 of the α1 CC domain, and D102 and P118 of the β1 H-NOX domain form hydrogen bonds with R258 of the β1 PAS domain (Figure 4E). To determine the function of these interfaces, alanine mutations were introduced to disrupt the interactions between them. We found that the D106A mutation in αF and the R258A mutation in βG of the β1 subunit both impaired NO activation (Figure 4F), suggesting that this interface plays an essential role in sGC activity in the NO-activated state.

### Structure of the sGC transducer module in the NO-activated state

The dramatically altered interactions between the CC domain and the sensor module lead to the conformational change of the transducer module (Figure 5). Strikingly, the linkers between αM and αN fold into α helical structures, which fuse the αM with αN into the 71 Å long αMN helixes (Figure 5A). Specifically, R420-K426 of the α1 CC domain and L355-Q358 of the β1 CC domain fold into α-helical structures (Figures 5B, S5, and S6). As a result, the transducer module switches from a highly bent conformation in the inactive state to a long, continuous coiled-coil structure in the NO-activated state (Figure 5B). The folding of the αM-αN loops results in a decrease in the exchangeability of the main chain hydrogens due to their hydrogen bonding in α helixes. This is consistent with previous hydrogen-deuterium exchange mass spectrum results that showed that the αM-αN loops had a significantly slower exchange rate upon NO activation (Underbakke et al., 2014). To determine the functional importance of this bending-straightening conformational change, we mutated residues in the αM-αN linker to either prolines or alanines. Prolines generate kinks in helical structures due to their incapability to form hydrogen bonds on the main chain. Therefore, proline mutations should destabilize the helical structures of αMNs in the NO-activated state, and these proline mutants may favor the inactive conformation. Indeed, proline mutations of D423 in the α1 CC domain or G356 in the β1 CC domain rendered sGC unresponsive for NO activation (Figures 5B-C). By contrast, mutation of the same set of residues into alanines had no such effect, indicating that the continuous helical structures of the αMNs are essential for NO activation of sGC (Figure 5C).

**Figure 5.**
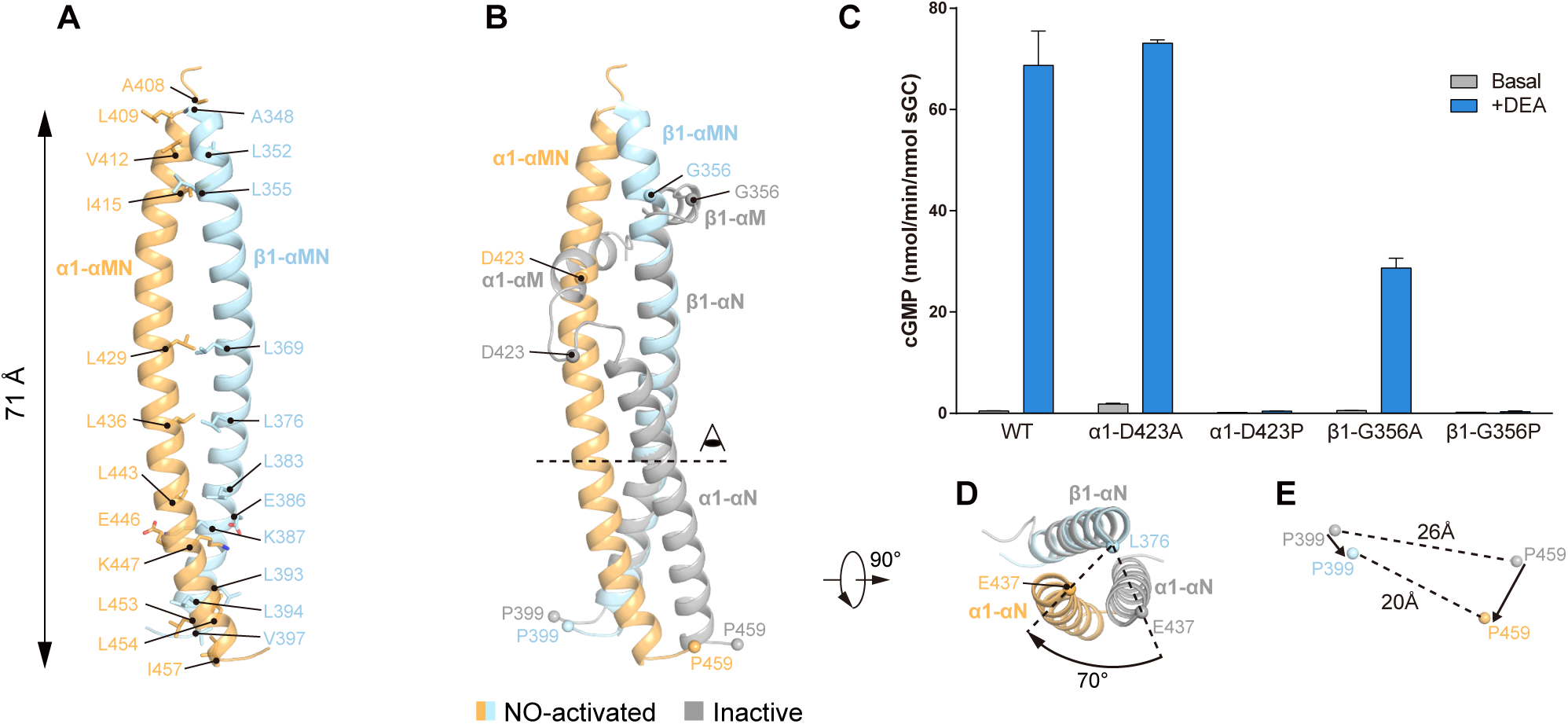
The structure of the sGC transducer module in the NO-activated state. (A) The transducer module in the NO-activated state, colored the same as in Figure 1A. (B) Structural comparison of the transducer module in the inactive state (gray) and the NO-activated state (colored the same as in Figure 1A) by aligning the αN helixes of their β1 subunits. The positions of Cα atoms of several key residues in the αM-αN linker are shown as spheres. (C) End point activity assay of the proline and alanine mutants in the αM-αN loop (Data are shown as means ± standard error, n = 3). (D) A 90° rotated top view of (B), beginning at the plane indicated with a dashed line and the point of view in (B). The arrow represents the rotation of the αN helix of the α1 subunit during activation. The angle between the Cα atoms of α1 E437 was measured using the Cα atom of β1 L376 as the vertex. (E) Positional displacement of the Cα atoms of β1 P399 and α1 P459. The Cα atoms in the inactive and NO-activated state are shown as gray spheres and colored spheres, respectively. The arrows denote the direction of change during activation.

In the NO-activated conformation, the interface between the α1 and β1 CC domain is drastically different from that observed in the inactive state. At the N-terminal part of the transducer module, A408, L409, V412, and I415 of α1 CC interact with A348, L352, and L355 of β1 CC (Figure 5A). At the middle of the transducer module, L429 and L436 of α1 CC interact with L369 and L376 of β1 CC (Figure 5A). At the C-terminal part of the transducer module, L443, L453, L454, and I457 of α1 CC make hydrophobic interactions with L383, L393, L394, and V397 of β1 CC (Figure 5A). In addition, E446 and K447 of α1 CC electrostatically interact with K387 and E386 of β1 CC (Figure 5A). Besides the overall “bending-straightening” movements of each CC domain, the αN helix of the α1 subunit rotates approximately 70° around the αN helix of the β1 subunit (Figure 5D). The separation of the C-termini of the transducer module also decreases. The distance of the Cα atoms between P459 of the α1 subunit and P399 of the β1 subunit shrinks from 26 Å to 20 Å (Figure 5E).

### Structure of the sGC catalytic module in the NO-activated state

As a consequence of the conformational change in the transducer module, the N-termini of the catalytic domains of each subunit move closer towards each other (Figure 6A). This drives the structural reorganization of the entire catalytic module. The catalytic domain of the α1 subunit rotates 17° relative to the β1 catalytic domain (Figures 6A-B). These movements reorganize the catalytic center to facilitate the binding of the substrate GTP and the cofactor Mg^2+^ ions. In the map of the NO-activated state, we observe a strong density corresponding to the substrate analogue GMPCPP (Figure 6C). The catalytic domains of both subunits contribute to GMPCPP binding. αS, βK, βL, and βM of the β1 subunit form the guanosine binding pocket and R552 of the β1 subunit binds the α phosphate of GMPCPP. D530 and D486 of the α1 subunit coordinate the Mg^2+^ ions, which are essential for catalysis. Positively charged R574 of the α1 subunit and K593 of the β1 subunit bind the γ phosphate of GMPCPP (Figure 6C). By comparing the current structure with the active adenylate cyclase structure (PDB ID: 1CJU) (Tesmer et al., 1999) (Figure 6D), we found that the residues responsible for substrate binding and catalysis are at similar positions, indicating that the current sGC structure represents a catalytically competent conformation. Interestingly, we found that the C-terminus of the α1 catalytic domain folds back to bind between αQ of the α1 subunit and αO-βK of the β1 subunit in the NO-activated state (Figure S4E), although the exact C-terminal residues could not be explicitly modeled, probably due to their high flexibility.

**Figure 6.**
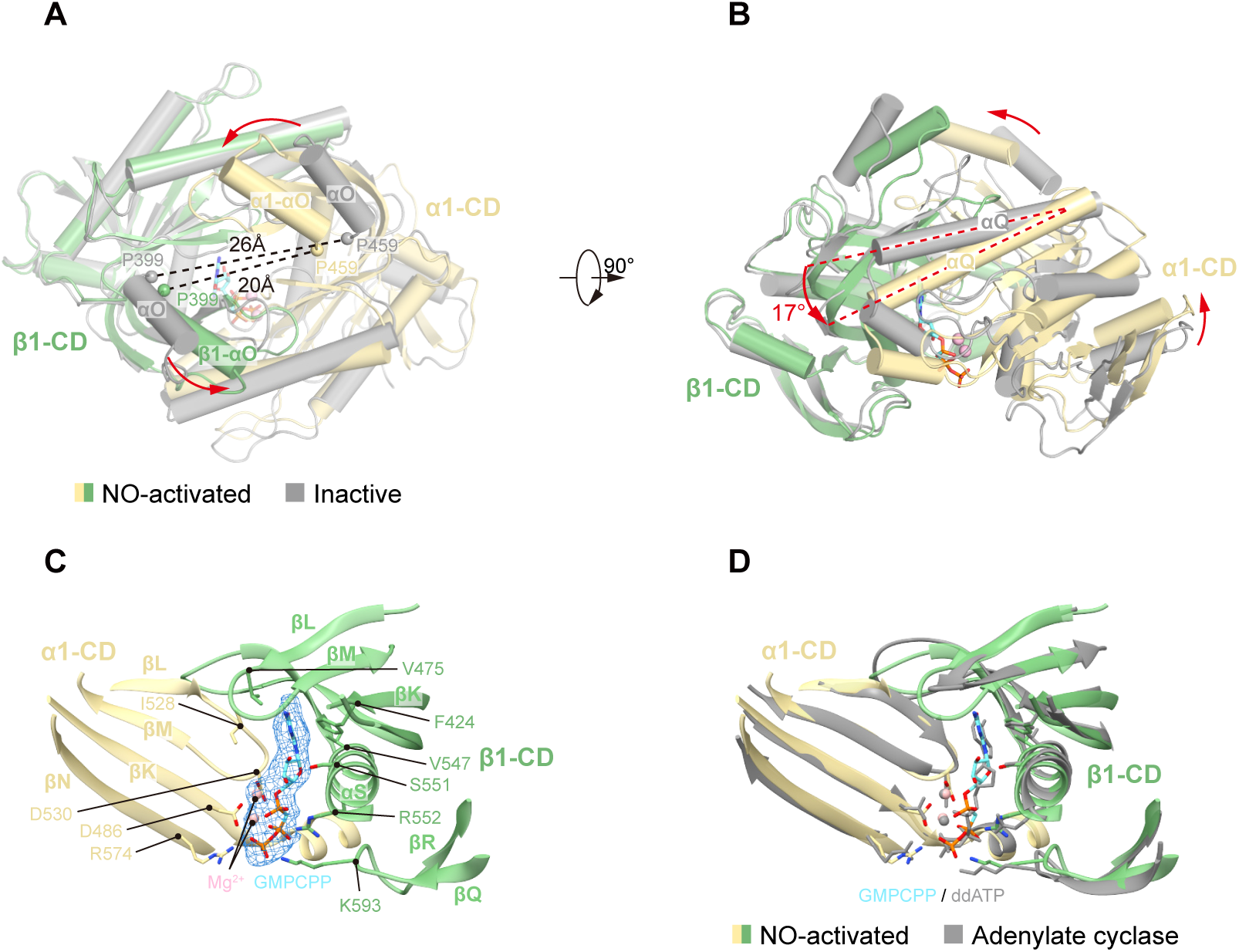
Structure of the sGC catalytic module in the NO-activated state. (A) Top view of the structural comparison of the catalytic module between the inactive state (gray) and the NO-activated state (colored). The colors of each domain in the NO-activated state are the same as in Figure 1A. The GMPCPP molecule is shown as sticks. The Cα atoms of α1 P459 and β1 P399 are shown as spheres. (B) A 90° rotated side view compared to (A). The angle between the αQ helixes in both the inactive state (gray) and the NO-activated state (colored) is shown. (C) The structure of the sGC catalytic core in the NO-activated state. GMPCPP is shown as cyan sticks and magnesium ions are shown as pink spheres. The densities of GMPCPP and magnesium are shown as blue mesh. The side chains of key residues that bind the substrate are shown as sticks. (D) Structural comparison of the catalytic core between the active adenylate cyclase (gray, PDB ID: 1CJU, chain A and B) and sGC in the NO-activated state (colored). The β1 subunit of sGC (green) is used for structural alignment.

### Structural mechanism of sGC activation

By analyzing the structures of individual sGC domains in both inactive and NO-activated states, we found that the structures of the scaffolding PAS dimer remain relatively unchanged among the different states, with a RMSD of 0.91 Å (Figure S7I). Therefore, we used the structures of the PAS dimer as a reference point to align and to compare the two full-length structures (Figures 7A-B). During activation, the α1 H-NOX domain has a small concomitant downward movement, while the interfaces between α1 H-NOX and its adjacent domains are largely maintained (Figure 4C). This suggested that the α1 H-NOX domain may play a role that is mainly structural instead of one involved in NO signal transduction (Video S5). This is in agreement with the activity assay result that the H-NOX domain of the α1 subunit is dispensable for NO activation (Figure 7C) (Koglin and Behrends, 2003). In contrast, the local conformational change of the β1 H-NOX domain upon NO binding drives the structural rearrangement of the sensor module (Figure 4C), which, along with previous functional data, suggested that the H-NOX domain of the β1 subunit plays an essential role in NO sensing (Derbyshire and Marletta, 2012). Indeed, complete removal of the β1 H-NOX domain rendered the sGC enzyme trapped in a relatively low activity state and unresponsive to NO activation (Figure 7C). This suggests that the β1 H-NOX domain in the NO-bound state is necessary to stabilize the sGC enzyme in an active conformation. Further supporting this conclusion, disruption of the interactions between β1 H-NOX and adjacent domains by mutation also diminished the NO activation (Figure 4F).

**Figure 7.**
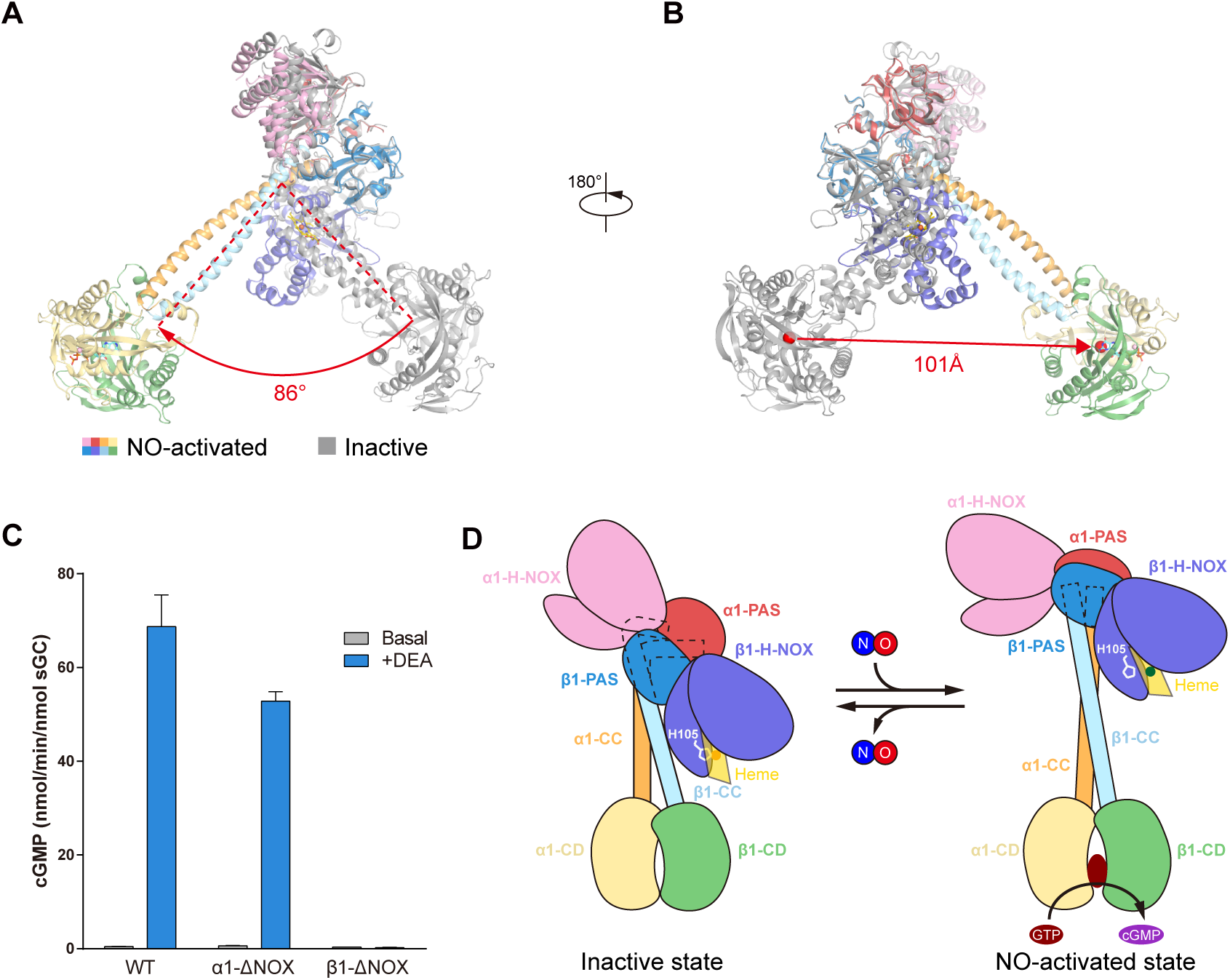
Overall structural rearrangement of sGC during NO activation. (A) The structure of sGC in the NO-activated state (colored the same as in Figure 1A) is overlaid onto the structure of sGC in the inactive state (gray) by superimposing the PAS domain dimer. The angle between the αN helixes of their β1 subunits is shown below. (B) A 180° rotated view compared to (A). The positional displacement of the centers of mass (red spheres) of the catalytic modules is shown below. (C) End point activity assay of the H-NOX domain deletion mutants (Data are shown as means ± standard error, n = 3). (D) Cartoon model of the conformational changes during sGC activation. The colors of each individual domain are the same as Figure 1A.

The structural changes in the sensor module upon NO binding trigger the bending-straightening conformational switch of the transducer module. As a result, the distal catalytic module rotates 86° in a swing-like manner and is spatially displaced by 101 Å (Figures 7A-B). Inhibition of the straightening by proline mutations abolishes the NO activation (Figure 5C), which suggests that the conformational change in the transducer module is essential for the activation of the catalytic module. Furthermore, we found that the catalytic module changes from a substrate binding incompatible conformation to a catalytically competent conformation (Figures 6A-B), which explained how the binding of NO decreases the *K*_m (GTP)_ and increases the *k*_cat_ of sGC (Denninger et al., 2000). However, the information flow in reverse direction, from the catalytic module to the sensor module, was implicated by previous functional data described in the literatures. Previous studies have reported that GTP binding to the catalytic module can stabilize the initial five-coordinate sGC-NO complex (Tsai et al., 2011). Moreover, ATP and GTP binding to the catalytic module slows NO dissociation (Cary et al., 2005). As a result, the catalytic module may act as a sensor for intracellular nucleotides and affect the sensitivity of sGC to NO (Ruiz-Stewart et al., 2004). We speculate that the nucleotide binding events in the catalytic module can alter its local conformation and send the signal via the middle transducer module to prime the sensor module and enhance NO sensitivity. Therefore, the transducer module acts as an allosteric structural coupler between the sensor module and the catalytic module to allow the bi-directional flow of information within the sGC molecule.

The structures of human α1β1 sGC presented here provide the first high-resolution view of the architecture and domain organization of this enzyme. Moreover, visualizing the sGC structures of multiple states, together with accompanying biochemical analysis, allowed us to characterize its allosteric activation mechanism (Figure 7D),. Our work not only paves the way towards a better mechanistic understanding of sGC activity, but also provides a template for structure-based drug design targeting related human diseases.

## Acknowledgments

We thank all of Chen Lab members for kindly help. We also thank Jiahuai Han for sharing the human sGC cDNAs and David B. Morton for sharing drosophila and mouse sGC cDNA. Cryo-EM data collection was supported by Electron microscopy laboratory and Cryo-EM platform of Peking University with the assistance of Xuemei Li, Zhenxi Guo, Bo Shao, and Xia Pei. Part of the structural computation was also performed on the Computing Platform of the Center for Life Science and High-performance Computing Platform of Peking University. The work is supported by grants from the Ministry of Science and Technology of China (National Key R&D Program of China, 2016YFA0502004 to L.C.) and National Natural Science Foundation of China (31622021 and 31521062, 31870833 to L.C.) and Young Thousand Talents Program of China to L.C. and the China Postdoctoral Science Foundation (2016M600856 and 2017T100014 to J.-X.W.). J.-X. W. is supported by the Boya Postdoctoral Fellowship of Peking University.

## Author Contributions

Lei Chen initiated the project and screened expression constructs. Rui Liu purified protein and prepared the cryo-EM sample. Yunlu Kang and Rui Liu collected the cryo-EM data with the help of Jing-Xiang Wu. Yunlu Kang processed the cryo-EM data with the help of Lei Chen. Lei Chen built and refined the atomic model. Yunlu Kang finished the enzymatic activity assay. Rui Liu finished the disulfide bond cross-linking experiment. All authors contributed to the manuscript preparation.

## Declaration of Interests

The authors declare no competing interests.

---

Video S1. Flexibility of sGC in the inactive state along the first eigenvector.

Video S2. Flexibility of sGC in the NO-activated state along the first eigenvector.

Video S3. Structure of sGC in the inactive state.

Video S4. Structure of sGC in the NO-activated state.

Video S5. Conformational changes of sGC during NO activation.

## Star Methods

### Cell Culture

HEK293F suspension cells were cultured in Freestyle 293 medium (Thermo Fisher Scientific) or SMM 293-TI medium (Sino Biological) supplemented with 1% FBS at 37°C with 6% CO _2_ and 70% humidity. It is reported that HEK293F is a female cell line. Sf9 insect cells were cultured in SIM SF (Sino Biological) at 27°C.

### Protein Expression and Purification

cDNA of *Drosophila melanogaster* (Vermehren-Schmaedick et al., 2010), mouse, and human sGC were cloned into a modified BacMam expression vector (Goehring et al., 2014; Li et al., 2017) and transfected into 293F cells for screening by fluorescence-detection size-exclusion chromatography (FSEC) (Kawate and Gouaux, 2006) on a Superose 6 increase 5/150 GL. The combination of C-terminal GFP-tagged human α1 and non-tagged β1 yielded a stable heterodimer. The two coding sequences were transformed into the pFastbac dual vector and expression was driven by p10 or polyhedrin promoters. The corresponding baculovirus was generated using the Bac-to-Bac system.

Sf9 insect cells at a density of 4 × 10^6^/mL in SIM SF medium were infected with the baculovirus and cultured at 27°C in a shaker for 72 h before harvesting and storage at −80°C. Cells corresponding to 500 ml culture were thawed and resuspended with 20 mL lysis buffer (50 mM Tris pH 8.0 at 4°C, 150 mM NaCl) containing 1 μg/mL aprotinin, 1 μg/mL pepstatin, 1 μg/mL leupeptin, 1 mM phenylmethanesulfonylfluoride, 2 mM dithiothreitol (DTT), and 1 mM EDTA. Cells were broken by sonication in 5 s intervals followed by a 5 s pause at 50% output for 20 min. Unbroken cells, cell debris, and membranes were removed by ultracentrifugation at 40,000 rpm for 1 h at 4°C using a Ti45 rotor (Beckman). Excess amount of purified glutathione S transferase-tagged GFP-nanobody (Tang et al., 2018) was added to the supernatant and incubated at 4°C for 10 min with rotation. Samples were then loaded onto 4 mL Glutathione Sepharose 4B columns (GE Healthcare) and washed with TBS buffer (20 mM Tris, pH 8.0, 150 mM NaCl) containing 1 mM dithiothreitol at 4°C. Protein was eluted with elution buffer (50mM Tris, pH 8.0, 10 mM reduced glutathione, 1 mM dithiothreitol) at room temperature. The eluate was diluted with buffer A (20 mM Tris, pH 8.0 at 4°C) to a conductance lower than 5000 Ω^-1^ and loaded onto a 1 mL HiTrap Q HP (GE Healthcare). The protein was eluted with buffer B (20 mM Tris, pH 8.0, 500 mM NaCl) at 4°C in a linear gradient using the AKTA pure system (GE Healthcare). The peak fractions containing sGC were pooled and incubated with prescission protease overnight to cleave the tag off from the protein. The digested protein was further purified by Superdex 200 increase (GE Healthcare) running in buffer containing 20 mM HEPES, 50 mM NaCl and 2 mM Tris (2-carboxyethyl) phosphine (TCEP). The peak fractions containing the sGC protein were pooled and concentrated. UV-Vis spectrum is measured using a spectrometer (Pultton) in the cuvette mode.

### Activity Assay

The protein used for cryo-EM sample preparation was diluted with 20 mM triethanolamine (pH 7.6), 300 mM NaCl, 1 mM dithiothreitol and subjected to activity assay as described below. For the heme-oxidized sample, the protein was diluted with 20 mM triethanolamine (pH 7.6), 300 mM NaCl and preincubated with 20 μM NS2028 at 25°C for 30 min and then added dithiothreitol to the final concentration of 1mM for activity assay. To generate the sGC mutant protein for activity assay, the coding sequences of the α1 subunit with a C-terminal GFP-strep tag and the β1 subunit were cloned into pFastBac1 expression vectors, respectively. To generate α1-ΔNOX and β1-ΔNOX constructs, the N-terminal 273 amino acids of the α1 subunit and 200 amino acids of the β1 subunit were removed, respectively. The constructs carrying desired point mutations were generated by Quick Change method and corresponding baculoviruses were generated by Bac-to-Bac system. Sf9 insect cells at the density of 4 × 10^6^/ml in SIM SF medium were infected with baculovirus and cultured in a shaker at 27°C for additional 72 h before harvest. Cells were resuspended with buffer containing 50 mM Tris (pH 8.0 at 4°C), 150 mM NaCl, 1 μg/ml aprotinin, 1 μg/ml pepstatin, 1 μg/ml leupeptin, 1 mM phenylmethanesulfonylfluoride and 1mM dithiothreitol, and broken by passing through the syringe needle with 0.45 μm inner diameter for 6 times. Cell debris and membrane were removed by ultracentrifugation at 40,000 rpm for 30min at 4°C using TLA55 rotor (Beckman). The supernatants were loaded onto Streptactin Beads 4 FF (Smart-Lifesciences) and washed with buffer containing 20 mM Tris (pH 8.0 at 4°C), 150 mM NaCl and 1 mM dithiothreitol. The protein was eluted with buffer containing 50 mM Tris (pH 8.5 at 4°C), 50 mM NaCl, 10 mM D-desthiobiotin and 1 mM dithiothreitol. The eluates were diluted with equal volume of 20 mM triethanolamine (pH 7.6) and loaded onto Q Beads 6 FF (Smart-Lifesciences) by gravity and washed with buffer containing 20 mM triethanolamine (pH 7.6), 150 mM NaCl and 1 mM dithiothreitol. The protein was eluted by 20 mM triethanolamine (pH 7.6), 300 mM NaCl and 1 mM dithiothreitol. The protein concentrations of various GFP-tagged sGC mutants were determined by comparing their GFP fluorescence signal to that of a purified GFP-tagged sGC standard on fluorescence-detection size-exclusion chromatography (FSEC) (Kawate and Gouaux, 2006). The activity assay mixture contained 10 nM sGC, 60 mM triethanolamine (pH 7.6), 150 mM NaCl, 0.5 mM dithiothreitol, 5 mM MgCl_2_, 200 µM GTP with or without 200 µM DEA NONOate (Cayman Chemical) in a final volume of 20 μl. The assay mixture was incubated at 25°C for 10 min and stopped by adding 80 μl 125 mM Zn(OAc)_2_ and 100 μl 125 mM Na_2_CO_3_. The GTP-ZnCO_3_ precipitation was removed by centrifugation at 17,000 × g for 5min and the supernatant was used for cGMP quantification, which was done by using the Cyclic GMP ELISA Kit (Cayman Chemical) according to the instruction. Each assay was independently repeated for at least 3 times.

### EM sample preparation

Purified sGC was concentrated and supplemented with the ligands before freezing. Protein samples were loaded onto glow-discharged Quantifoil 0.6/1 holey carbon gold grids and plunged into liquid ethane by Vitrobot Mark IV (Thermo Fisher Scientific).

### Disulfide bond cross-linking

To generate the cys-less sGC construct (sGC^CL^), the cys-rich N-terminal 63 amino acids of α1 subunit were removed. Additional mutations of C176A, C239A, C669S, C455Y, and C460G were done on α1 subunit and C292N were done on β1 subunit. The coding sequences of α1^CL^ with C-terminal GFP-strep tag and β1^CL^ subunit without tags were cloned into modified BacMam expression vectors (Goehring et al., 2014; Li et al., 2017) respectively. Then specific amino acids were mutated into cysteines by Quick Change method. Cysteine mutants were transfected into HEK293F cells with 3 μg/ml polyethylenimines (PEIs) (Polysciences) at the density of 2.0 × 10^6^/mL. Cells were harvested 72 h post transfection and broken by passing the syringe needle with 0.45 µm inner diameter for 10 times. Unbroken cells and large debris were removed by centrifuged at 14,800 rpm at 4°C for 10 min. sGC protein were purified from supernatants by Streptactin Beads 4FF resin (Smart-Lifesciences). Protein samples were cross-linked on ice for 30 min by adding Cu(II) (1,10-phenanthroline)_3_ to the final concentration of 0.1 mM to promote disulfide bond formation. Protein samples were subjected to 4-15% gradient SDS-PAGE (BeyoGel) for separation either in non-reducing condition or reducing condition (in the presence of 100 mM DTT). The fluorescence was detected in ChemiDoc MP (Bio-Rad) fluorescence imaging system.

### Cryo-EM Data Acquisition

Cryo-grids were screened on a Talos Arctica electron microscope (Thermo Fisher Scientific) operated at 200 kV using a Ceta 16M camera (Thermo Fisher Scientific). The screened grids were transferred to a Titan Krios electron microscope (Thermo Fisher Scientific) operated at 300 kV with an energy filter set to a slit width of 20 eV. Images were recorded using a K2 Summit direct electron camera (Thermo Fisher Scientific) in super-resolution mode at a nominal magnification of 130,000 ×, corresponding to a calibrated super-resolution pixel size of 0.5225 Å. The defocus range was set from −1.5 μm to −2 μm. Each image was acquired as a 7.68 s movie stack (32 frames) with a dose rate of 6.25 e^-^Å^-2^s^-1^, resulting in a total dose of about 48 e^-^Å^-2^. All data acquisition was done using SerialEM.

### Cryo-EM Data Processing

The data processing workflows are illustrated in Figures S1-4 and Table S1. Super-resolution movie stacks were motion-corrected, mag-distortion corrected, dose-weighted, and binned to a pixel size of 1.045 Å by MotionCor2 1.1.0 using 5 × 5 patches (Zheng et al., 2017). Contrast transfer function (CTF) parameters were estimated from non-dose-weighted micrographs using Gctf v1.06 (Zhang, 2016). Micrographs with ice or ethane contamination, empty carbon, and poor CTF fit (> 5 Å) were manually removed. All classification and reconstruction was performed with Relion 3.0 (Zivanov et al., 2018) unless otherwise stated. Particles were picked using Gautomatch (developed by Kai Zhang) and subjected to reference-free 2D classification to remove bad particles. Initial models were generated by cryoSPARC (Punjani et al., 2017) using the selected particles from 2D classification. The particles were further subjected to 3D classification to remove bad particles using the initial model, which was low-passed filtered to 30 Å as the reference. The particles selected from good 3D classes were re-centered and re-extracted, and their local CTF parameters were individually determined using Gctf v1.06 (Zhang, 2016). These particles were imported into cisTEM (Grant et al., 2018) and subjected to 3D classification with auto-masking. The particles from the best 3D classes calculated by cisTEM were exported into Relion 3.0 and subjected to 3D auto-refinement to generate the consensus map. Two soft masks that cover the N lobe and C lobe were generated from the consensus map, which was edited manually in UCSF Chimera using the volume eraser tool (Pettersen et al., 2004). 3D multi-body refinements (Nakane et al., 2018) were performed using the two soft masks of the lobes and the parameters determined from previous 3D auto-refinement. The motions of the bodies were analyzed by relion_flex_analyse in Relion 3.0. The two half-maps of each lobe generated by 3D multi-body refinement were subjected to post-processing in Relion 3.0. The masked and sharpened maps of each lobe were aligned to the consensus map using UCSF Chimera and summed to generate the composite map for visualization and interpretation. All of the resolution estimations were based on the Fourier shell correlation (FSC) of 0.143 cutoff after correction of the masking effect (Chen et al., 2013). B-factors used for map sharpening were automatically estimated by the post-processing procedure in Relion 3.0.

### Model building

The position of β1 H-NOX domain was firstly identified according to its distinguishable heme group density. Other domains were assigned by the domain-domain linkers that are visible in the post-processed map (Figure S2G). The homology models of individual H-NOX, PAS and catalytic domains were generated by Phyre2 server (Kelley et al., 2015) based on the structures of human β1 H-NOX (PDB:5MNW), *Manduca sexta* α PAS domain (PDB: 4GJ4) (Purohit et al., 2013) and human α1, β1 catalytic domain heterodimer (PDB:3UVJ) (Allerston et al., 2013). The models were placed into the corresponding composite maps using UCSF chimera (Pettersen et al., 2004) and manually rebuilt in Coot (Emsley et al., 2010). The composite maps were then converted into mtz files and the models were further refined by Phenix in reciprocal space (Adams et al., 2010) and Coot in real space. During model building, we found the structure of catalytic module of heme-oxidized state and the heme-unliganded state are essentially the same, but we observed a positive difference density around the βK-αP loop of α1 catalytic domain in the heme-oxidized state sample (Figure S2H). During the sample preparation of the heme-oxidized state, we supplemented oxidizing reagent NS2028, substrate GTPγS and cofactor Mg^2+^ ions into the sGC protein. Therefore, based on the local chemical environment, this positive density might represent Mg^2+^ ions together with highly negatively charged phosphate groups that possibly came from the decomposition of the GTPγS molecule. However, this putative phosphate groups were not modeled.

### Quantification and statistical analysis

Global resolution estimations of cryo-EM density maps are based on the 0.143 Fourier Shell Correlation criterion (Rosenthal and Henderson, 2003). The local resolution was estimated using relion_postprocess in Relion 3.0 (Zivanov et al., 2018). The number of technical replicates (N) and the relevant statistical parameters for each experiment (such as mean or standard error) are described in the figure legends. No statistical methods were used to pre-determine sample sizes.

### Data availability

Cryo-EM maps and atomic coordinates of the heme-unliganded, heme-oxidized, NO-activated and β1 H105C mutant sGC have been deposited in the EMDB and PDB under the ID codes EMDB: EMD-9883, EMDB9884, EMDB9885, EMDB-9886 and PDB: 6JT0, 6JT1, 6JT2, respectively.

---

**Figure S1.**
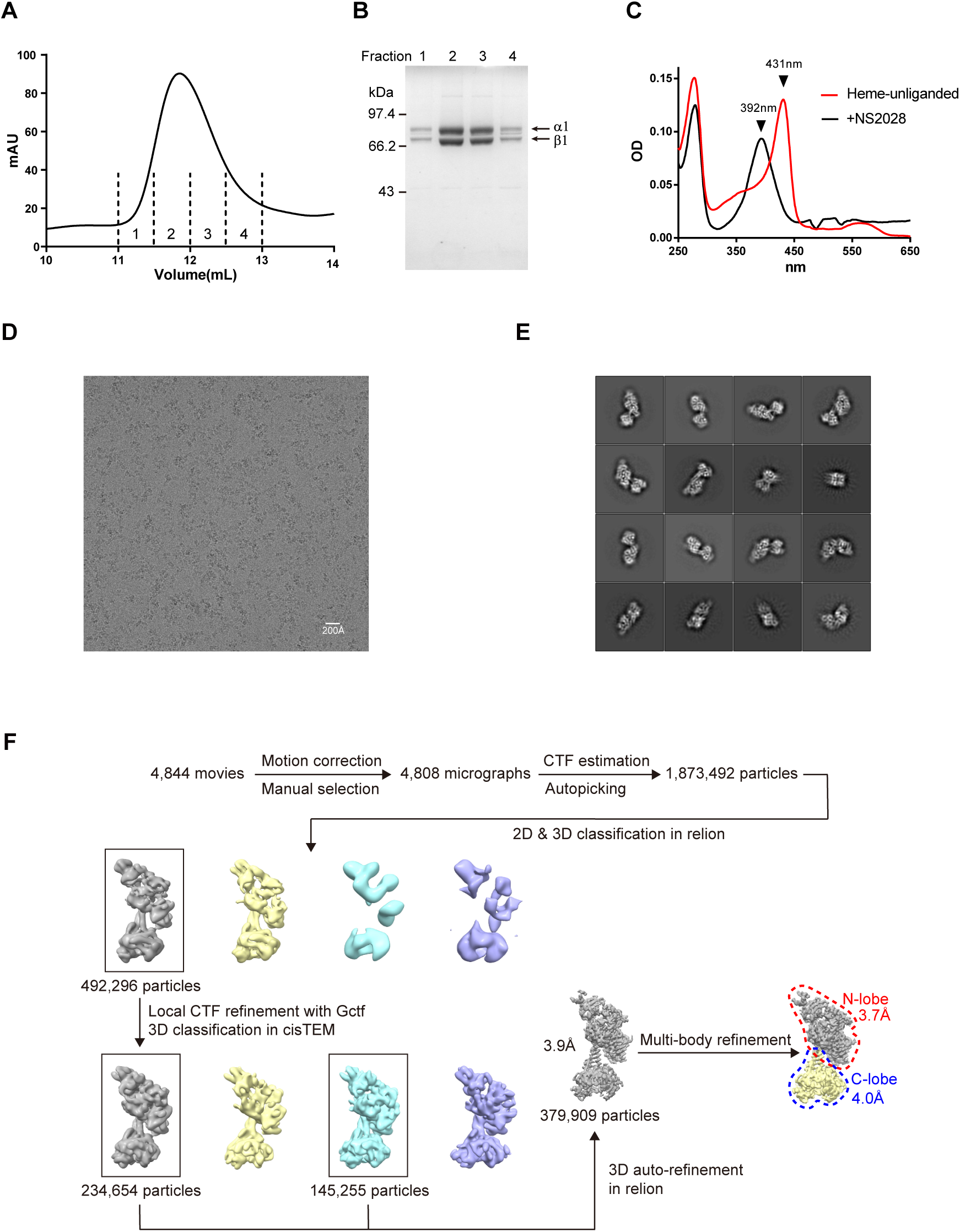
Biochemical characterization of the human α1β1 sGC heterodimer protein and single particle cryo-EM data processing procedure of sGC in the inactive (heme-oxidized) state. Related to Figure 1. (A) Size-exclusion chromatography of sGC on a superdex 200 column. The fractions indicated by dashed lines were pooled for cryo-EM sample preparation. (B) SDS-PAGE of the size-exclusion chromatography fractions labeled in (A). The positions of the α1 and β1 subunits are denoted with arrows. (C) UV-Vis spectrum of purified sGC before (red) and after (black) NS2028 treatment. The positions of the Soret peaks are indicated by arrows. (D) A representative raw micrograph of sGC in the inactive (heme-oxidized) state. (E) Representative 2D class averages of sGC in the inactive (heme-oxidized) state. (F) The cryo-EM data processing workflow of sGC in the inactive (heme-oxidized) state.

**Figure S2.**
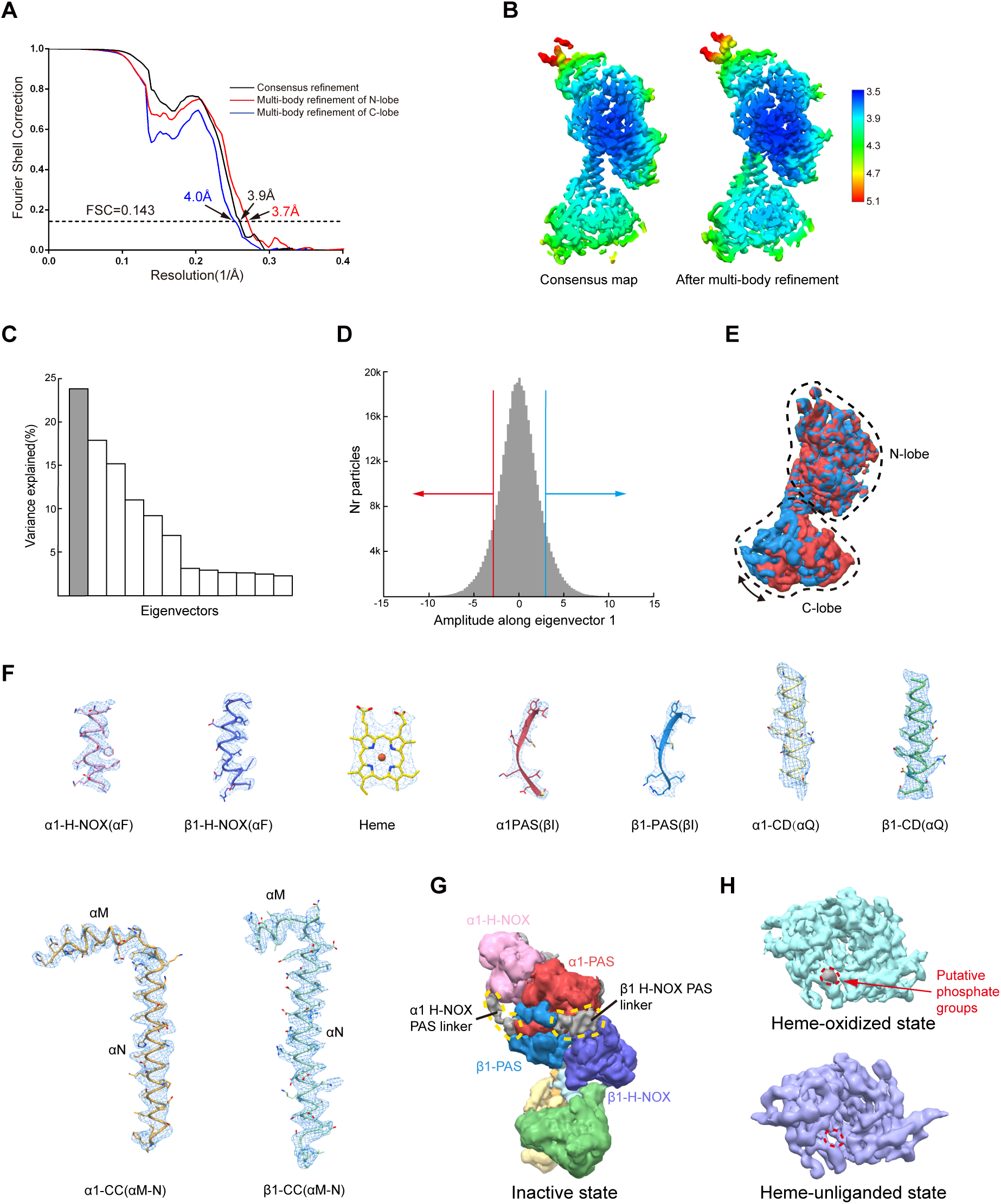
Conformational heterogeneity and local density quality of sGC in the inactive (heme-oxidized) state. Related to Figure 1. (A) Gold-standard Fourier shell correlation (FSC) curves of the heme-oxidized sGC after correction for the masking effects. Resolution estimations were based on the criterion of the FSC 0.143 cutoff. (B) Local resolution distribution of the composite map of sGC in the inactive (heme-oxidized) state. (C) Histogram of the eigenvectors that contribute to the variance. The top eigenvector is highlighted in gray. (D) Histogram of the amplitudes along the top eigenvectors shows monomodal distribution. Particle populations with amplitudes less than −3 or greater than 3 are indicated as red and blue arrows, respectively. (E) 4 Å low-passed filtered maps reconstructed from particles indicated as red and blue arrows in (D). N-lobes were used for alignment. (F) Representative cryo-EM densities of fragments from each individual domain. (G) The cryo-EM map of the sGC in the heme-oxidized state. The putative linkers between H-NOX domain and PAS domain were shown in gray. B-factor of the map was adjusted to −100 Å2 in post-processing procedure to visualize features with high flexibility. (H) The cryo-EM map of the catalytic module in the heme-oxidized state (cyan) and the heme-unliganded state (purple). Density of the putative phosphate groups is shown in gray.

**Figure S3.**
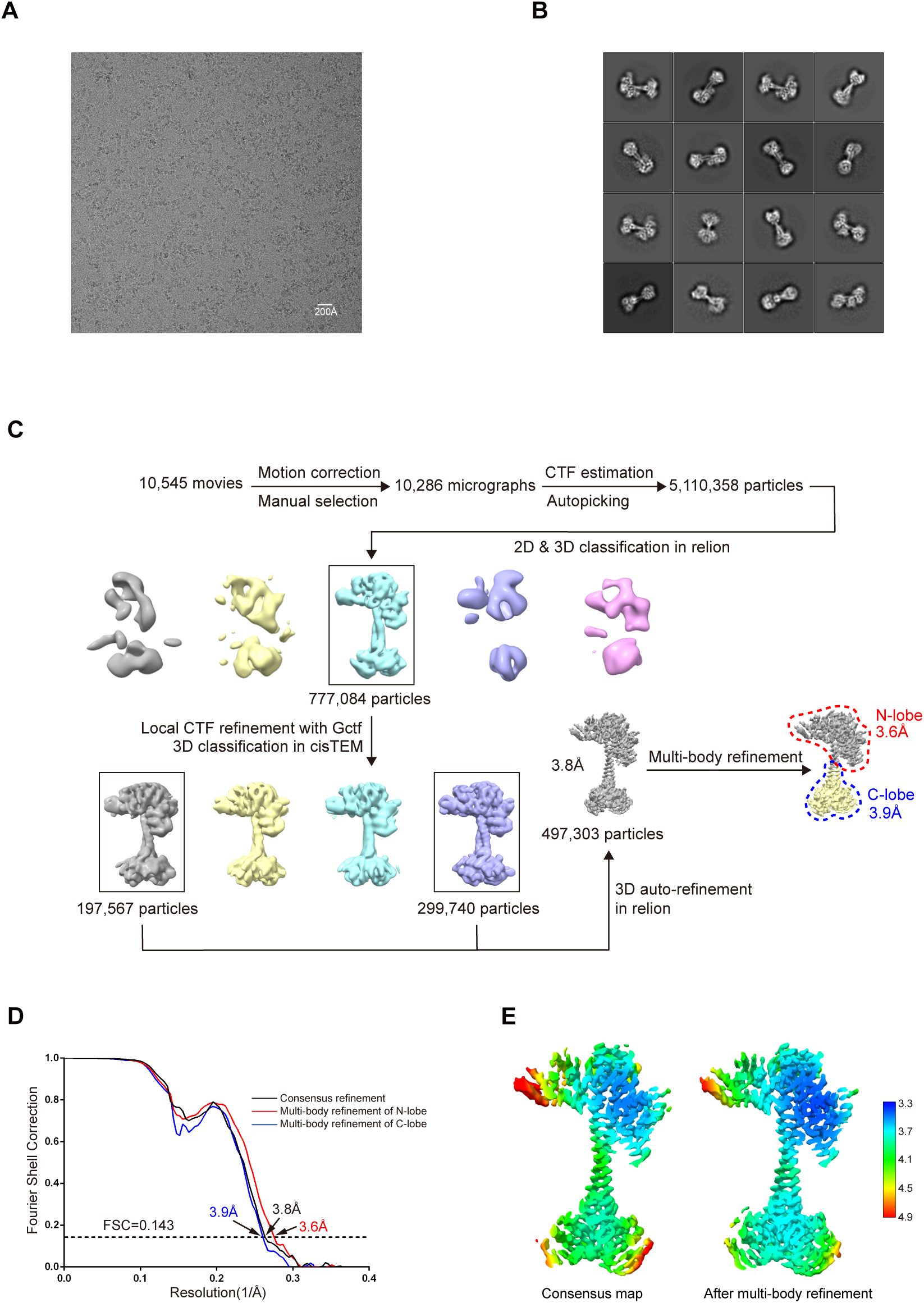
Single particle cryo-EM data processing procedure of the sGC sample in the NO-activated state. Related to Figure 3. (A) Representative raw micrograph of sGC in the NO-activated state. (B) Representative 2D class averages of sGC in the NO-activated state. (C) The cryo-EM data processing workflow of sGC in the NO-activated state. (D) Gold-standard FSC curves (after correction for the masking effects) of the NO-activated sGC. Resolution estimations were based on the criterion of FSC 0.143 cutoff. (E) Local resolution distribution of the composite map of sGC in the NO-activated state.

**Figure S4.**
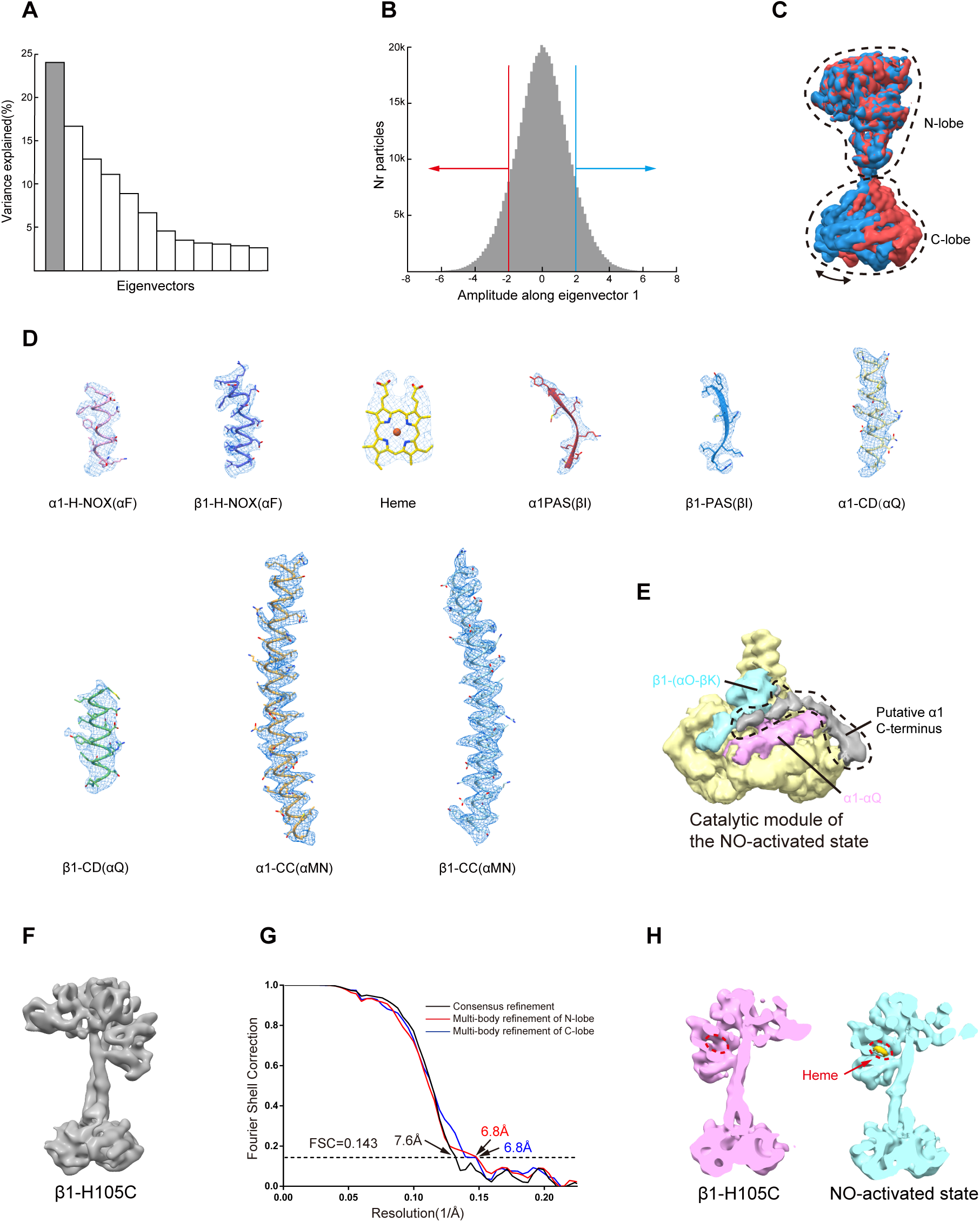
Conformational heterogeneity and local density quality of sGC in the NO-activated state. Related to Figure 3. (A) Histogram of the eigenvectors that contribute to the variance. The top eigenvector is highlighted in gray. (B) Histogram of the amplitudes along the top eigenvectors shows monomodal distribution. Particle populations with amplitudes less than −2 or greater than 2 are indicated as red and blue arrows, respectively. (C) 4 Å low-passed filtered maps reconstructed from particles indicated as red and blue arrows in (B). N-lobes were used for alignment. (D) Representative cryo-EM densities of fragments from each individual domain. (E) The cryo-EM map of the catalytic module in the NO-activated state (yellow). Putative density of the α1 C-terminus is shown in gray, the αQ of the α1 subunit is shown in pink, and the αO-βK fragment of the β1 subunit is shown in cyan. The B-factor of the map was adjusted to −100 Å2 in post-processing to visualize features with high flexibility. (F) The side view of the cryo-EM map of the β1 H105C mutant sGC. (G) Gold-standard FSC curves (after correction for the masking effects) of the β1 H105C mutant sGC. Resolution estimations were based on the criterion of FSC 0.143 cutoff. (H) The cryo-EM map of the β1 H105C mutant sGC (pink) and the NO-activated sGC (cyan). Density of the heme is shown in yellow. The map of NO-activated sGC was low-passed filtered to 6.8 Å.

**Figure S5.**
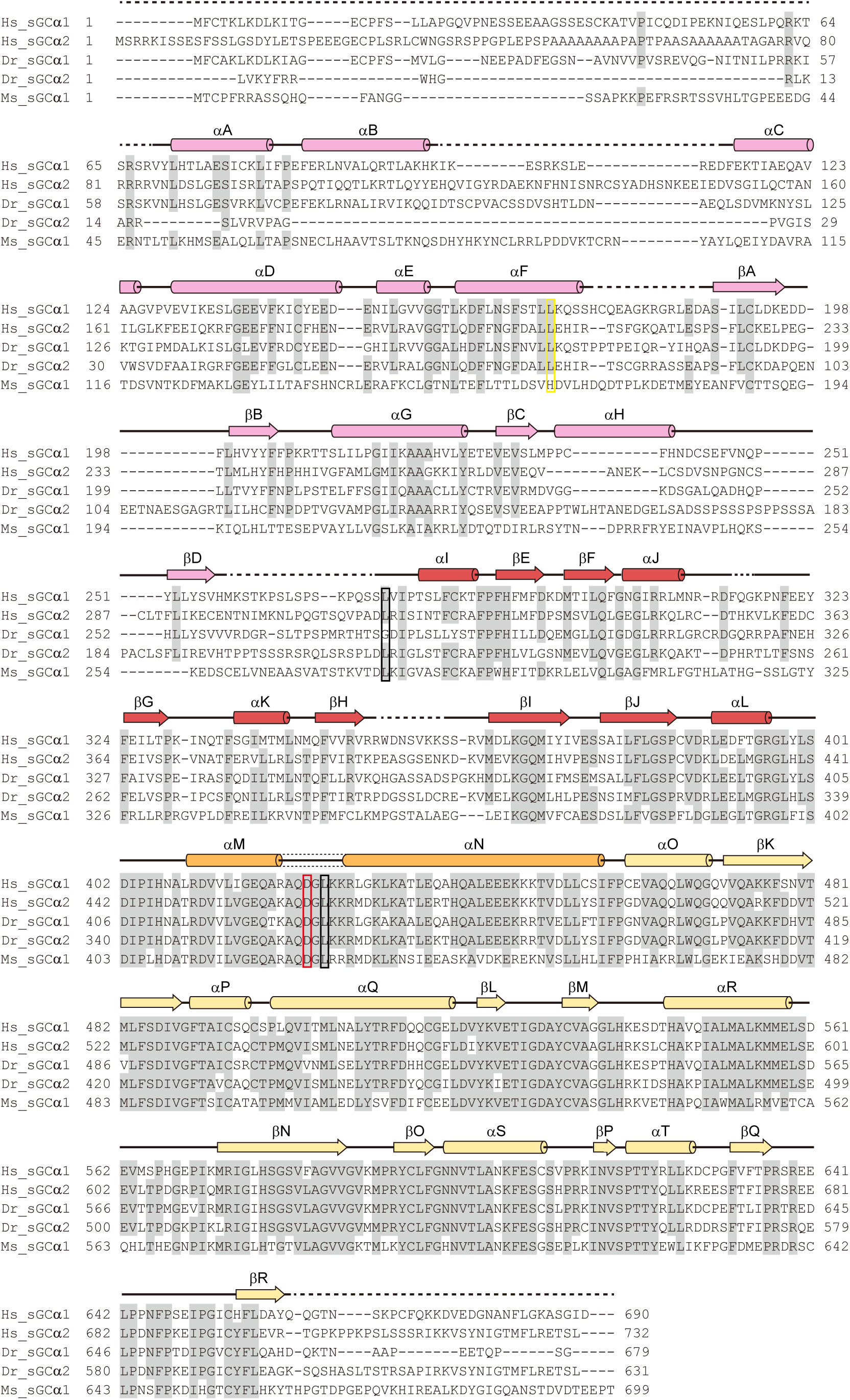
Sequence alignment of the sGC α subunit. Related to Figure 1. The sequences of the *Homo sapiens* α1 subunit, *Homo sapiens* α2 subunit, *Danio rerio* α1 subunit, *Danio rerio* α2 subunit, and *Manduca sexta* α1 subunit were aligned. Conserved residues are colored in gray. Residues that are mutated to cysteines for oxidative cross-linking are indicated with black box. Mutations for activity assay are indicated with red box. The residues corresponding to H105 in the β1 subunit are indicated with yellow box. Secondary structural elements are indicated as follows: β sheets by arrows, α helixes by cylinders, and loops by lines. Unmodeled residues are shown as dashed lines. The color of arrows and cylinders are the same as in Figure 1A.

**Figure S6.**
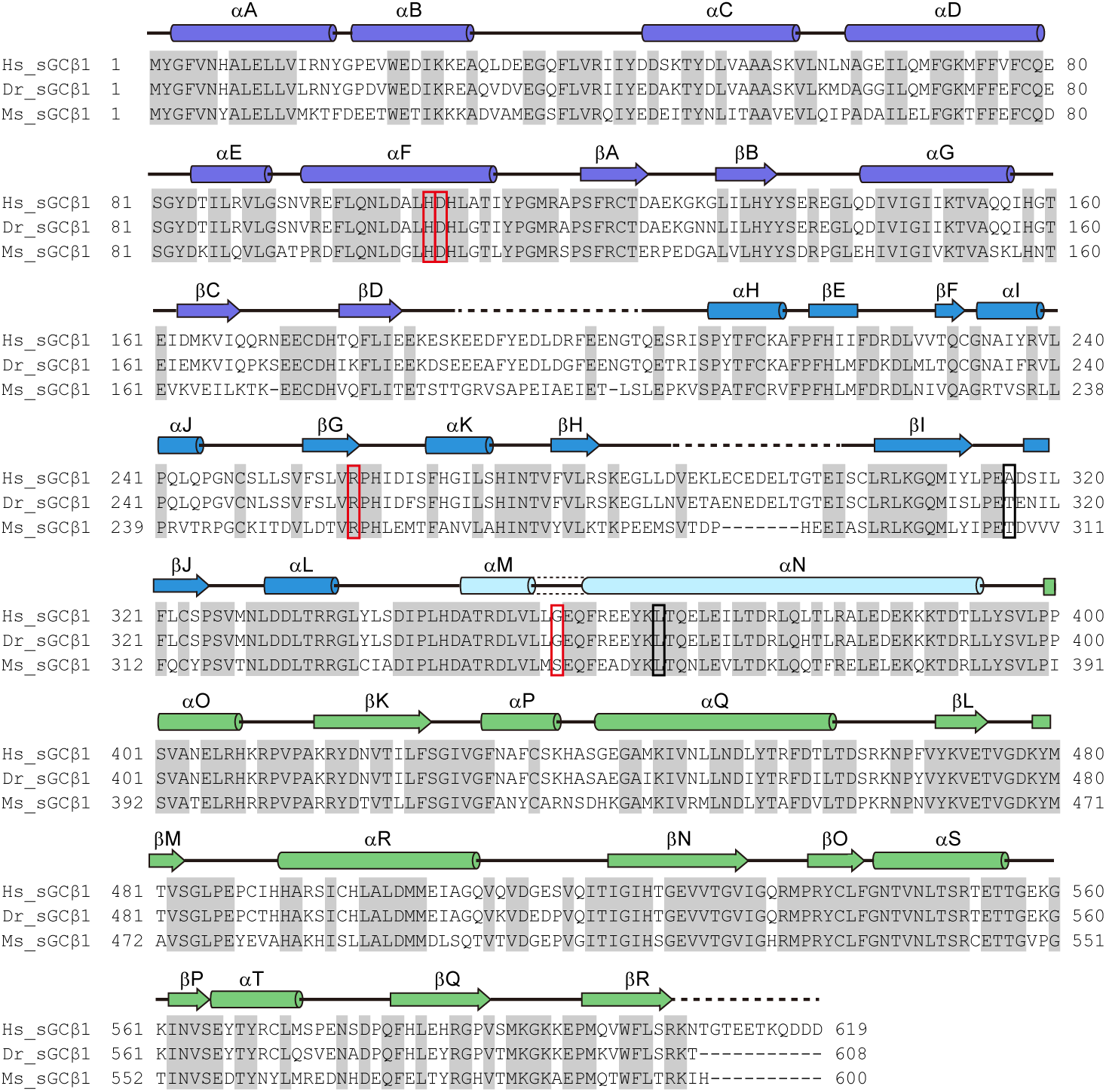
Sequence alignment of the sGC β1 subunit. Related to Figure 1. The sequences of the *Homo sapiens* β1 subunit, *Danio rerio* β1 subunit, and *Manduca sexta* β1 subunit were aligned. Conserved residues are colored in gray. Residues that are mutated to cysteines for oxidative cross-linking are indicated with black box. Mutations for activity assay are indicated with red box. Secondary structural elements are indicated as follows: β sheets by arrows, α helixes by cylinders, and loops by lines. Unmodeled residues are shown as dashed lines. The color of arrows and cylinders are the same as in Figure 1A.

**Figure S7.**
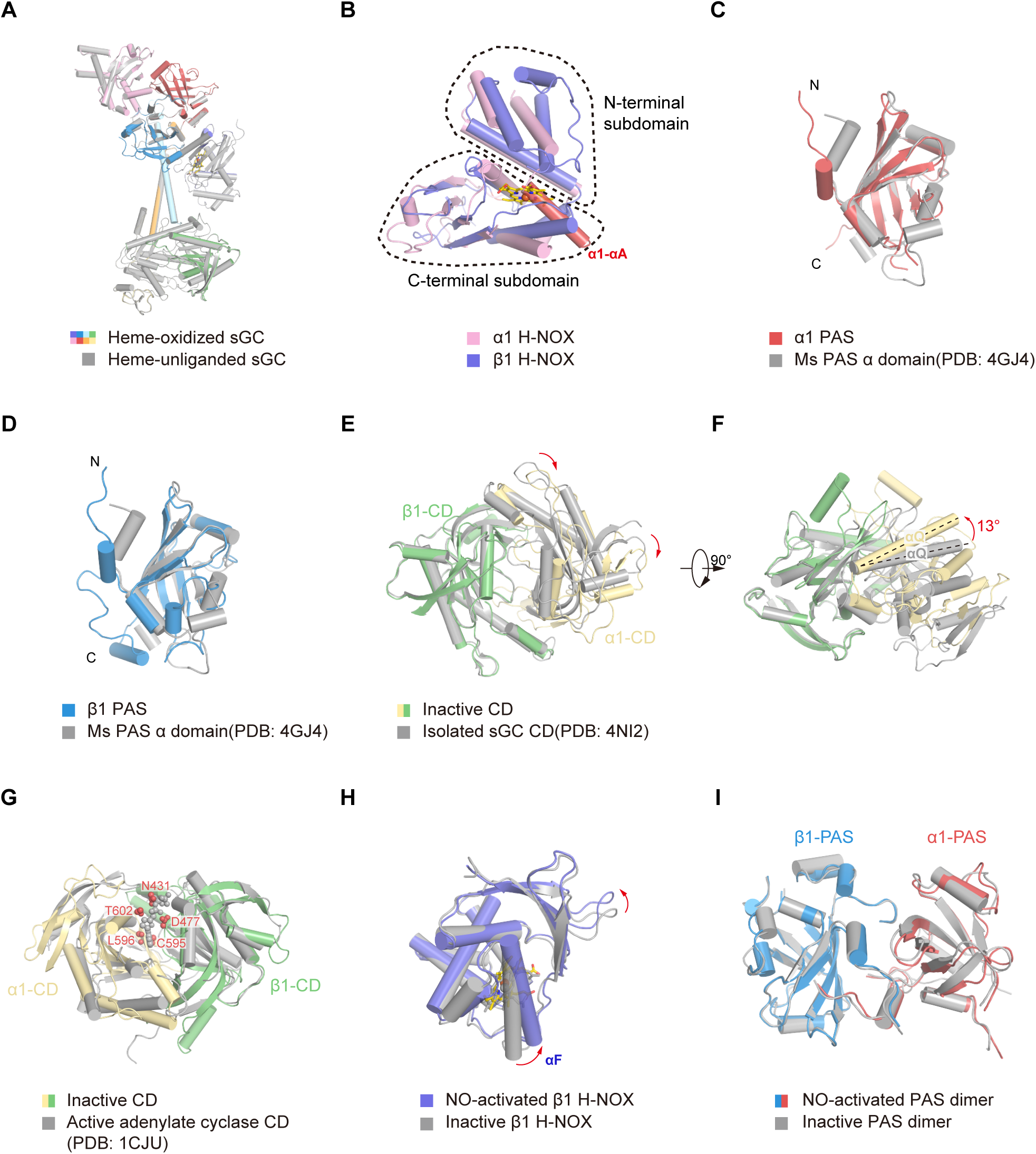
Structural comparisons of each domain. Related to Figure 2. (A) Structural comparison of the full-length human sGC between the heme-oxidized state (colored) and the heme-unliganded state (gray). (B) Structural comparison between the α1 H-NOX domain (pink) and the β1 H-NOX domain (blue) in the inactive state. The N-terminal αA helix of the α1 subunit that occupies the heme binding pocket is shown in red. The heme molecule of the β1 H-NOX domain is shown as a yellow stick. The approximate boundaries of N-terminal and C-terminal subdomains are indicated by dashed lines. (C) Structural comparison between the human α1 PAS domain (red) and the *Manduca sexta* Ms α PAS domain (gray, PDB ID:4GJ4). (D) Structural comparison between the human β1 PAS domain (blue) and the *Manduca sexta* Ms α PAS domain (gray, PDB ID:4GJ4). (E) Structural comparison between the catalytic module of the full-length sGC in the inactive state (colored) and the isolated catalytic domain heterodimer (gray, PDB ID: 4NI2). The β1 subunit was used for structural alignment. (F) 90° rotated view compared to (E). (G) Structural comparison between the catalytic module of the full-length sGC in the inactive state (colored) and the catalytic domain of the active adenylate cyclase (gray, PDB ID: 1CJU, chain A&B). The β1 subunit was used for structural alignment. The residues of sGC that are within 2.2 Å of the substrate are shown as red spheres. (H) Structural comparison between the NO-activated state (purple) and the inactive state (gray) of the human β1 H-NOX domain. The N-terminal subdomain was used for alignment and the movements are indicated as red arrows. (I) Structural comparison between the NO-activated state (colored) and the inactive state (gray) of the human α1 and β1 PAS heterodimer.)

**Supplementary Table. 1.**
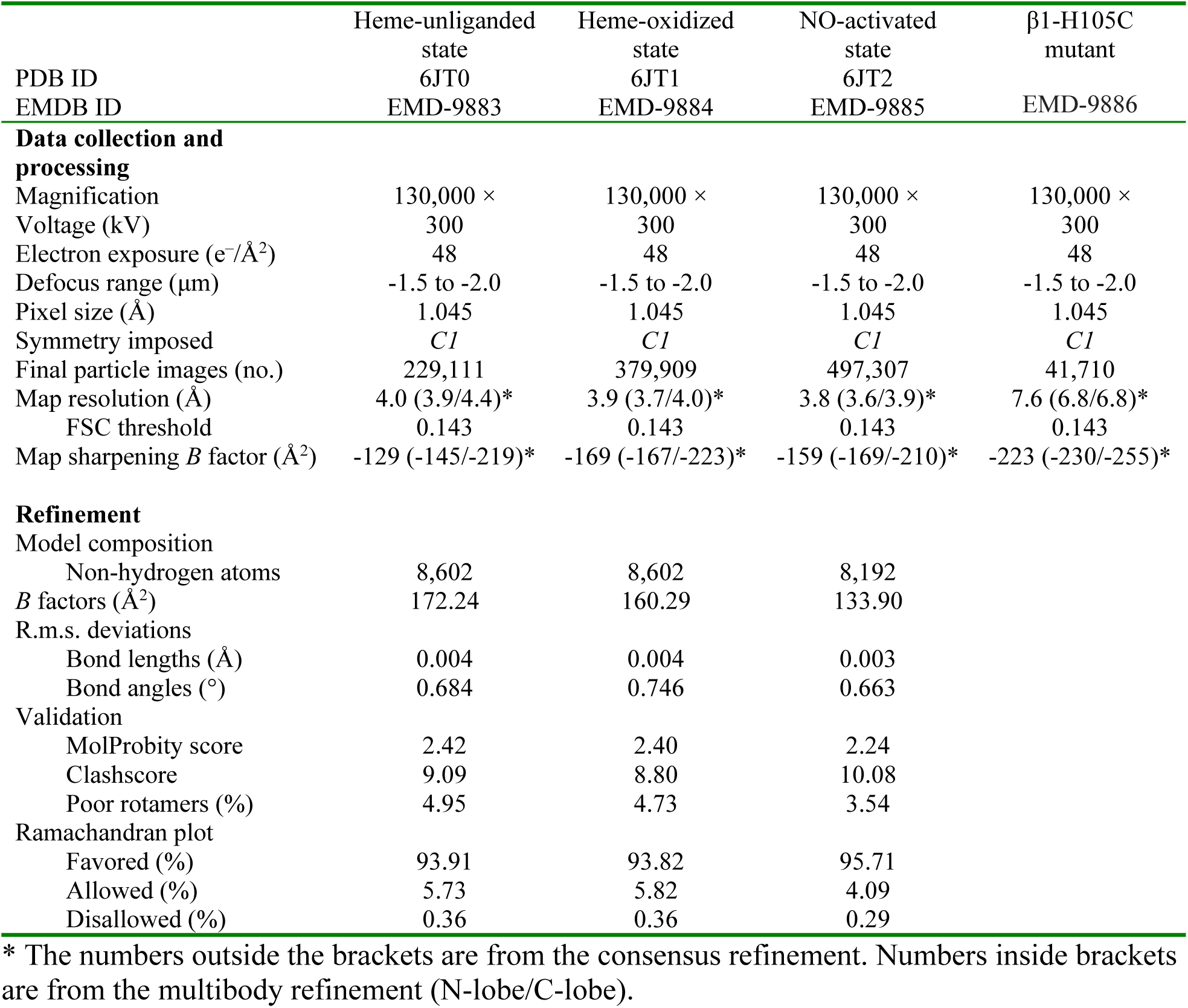
Cryo-EM data collection, refinement and validation statistics.

